# A new test suggests that balancing selection maintains hundreds of non-synonymous polymorphisms in the human genome

**DOI:** 10.1101/2021.02.08.430226

**Authors:** Vivak Soni, Michiel Vos, Adam Eyre-Walker

## Abstract

The role that balancing selection plays in the maintenance of genetic diversity remains unresolved. Here we introduce a new test, based on the McDonald-Kreitman test, in which the number of polymorphisms that are shared between populations is contrasted to those that are private at selected and neutral sites. We show that this simple test is robust to a variety of demographic changes, and that it can also give a direct estimate of the number of shared polymorphisms that are directly maintained by balancing selection. We apply our method to population genomic data from humans and conclude that more than a thousand non-synonymous polymorphisms are subject to balancing selection.

## Introduction

How genetic variation is maintained, either in the form of DNA sequence diversity or quantitative genetic variation, remains one of the central problems of population genetics. Balancing selection encapsulates several selective mechanisms that increase variability within a population. These include heterozygote advantage (also referred to as overdominance), frequency dependent selection, and selection that varies through space and time (Nielsen, 2005). However, although there are some clear examples of each type of selection (Allison, 1956; Nosil, et al., 2018), the overall role that balancing selection plays in maintaining genetic variation, either directly, or indirectly through linkage, remains unknown.

A number of methods have been developed to detect the signature of balancing selection (Hughes and Nei, 1988; Asthana et al. 2004; Bubb et al. 2006; Andres et al. 2009; Leffler et al. 2013; Degiorgio et al. 2014; Gao et al. 2015; Hunter-Zinck & Clark, 2015; Fijarczyk & Babik, 2015; Sheehan & Song, 2016; Siewert & Voight, 2017; Bitarello et al. 2018). Application of these methods have identified a number of loci subject to balancing selection, largely in the human genome, in which most of this research has taken place. However, these methods are generally quite complex to apply, often leveraging multiple population genetic signatures of balancing selection and many require simulations to determine the null distribution. Furthermore, they do not readily yield an estimate of the number of polymorphisms that are directly subject to balancing selection, as opposed to being in linkage disequilibrium. Here we introduce a method that is simple to apply and which generates a direct estimate of the number of polymorphisms subject to balancing selection.

One signature of balancing selection that has been utilised in several studies is the sharing of polymorphisms between species (Asthana et al. 2004; Leffler et al. 2013; Gao et al. 2015). If the species are sufficiently divergent that they are unlikely to share neutral polymorphisms, then shared genetic variation can be attributed to balancing selection. These studies have concluded that there are relatively few balanced polymorphisms that are shared between humans and chimpanzees (Asthana et al. 2004; Leffler et al. 2013). However, this test is likely to be weak because humans and chimpanzees diverged millions of years in the past and it is unlikely that any shared selection pressures will be maintained over that time period.

The major problem with approaches that consider the sharing of polymorphisms between species or populations is differentiating selectively maintained polymorphisms from neutral variation inherited from the common ancestor. This problem can be solved by comparing the number of shared polymorphisms at sites which are selected, to those that are neutral. We expect the number of shared polymorphisms at selected sites to be lower than at neutral sites because many mutations at selected sites are likely to be deleterious, and hence unlikely to be shared. However, we can estimate the proportion that are effectively neutral by considering the ratio of polymorphisms, which are private to one of the two populations or species, at selected versus neutral sites. Although the method can be applied to any group of neutral and selected sites that are interspersed with one another we will characterise it in terms of non-synonymous and synonymous sites. Let the numbers of polymorphisms that are shared between two populations or species be S_N_ and S_s_ at non-synonymous and synonymous sites respectively, and the numbers that are private to one of the populations be R_N_ and R_S_ respectively. Let us assume that synonymous mutations are neutral and non-synonymous mutations are either neutral or strongly deleterious. Then it is evident that 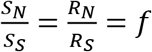, where *f* is the proportion of the non-synonymous mutations that are neutral. However, if there is balancing selection acting on some non-synonymous SNPs and this selection persists for some time such that the balanced polymorphisms are shared between populations then 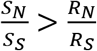. A simple test of balancing selection is therefore whether Z > 1 where

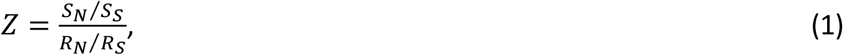

a simple corollary of the McDonald-Kreitman test for adaptive divergence between species (McDonald and Kreitman, 1991). It can be shown, under some simplifying assumptions in which synonymous mutations are neutral and non-synonymous mutations are strongly deleterious, neutral or subject to balancing selection, that an estimate of the proportion of non-synonymous mutations subject directly to balancing selection is 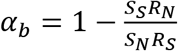 (see results section). In this analysis, we perform population genetic simulations to investigate whether the method can detect the signature of balancing selection and assess whether the method is robust to demographic change. Second, we apply the method to human population genetic data. We show that the method is robust and we estimate that substantial numbers of non-synonymous polymorphisms are maintained by balancing selection in humans.

## Results

### Simulations

We propose a new test for balancing selection in which the ratio of selected to neutral polymorphisms is compared between those that are shared between populations or species and those that are private to populations or species. To explore the properties of our method to detect balancing selection we ran a series of simulations in which an ancestral population splits to yield two descendent populations. We initially simulated loci under a simple stationary population size model where the ancestral population is duplicated to form two equally sized populations (equal to each other and the ancestral population). This is an unrealistic scenario, but it has the advantage that it involves no demographic change in the transition from ancestral to descendent populations. We assume that synonymous mutations are neutral and we explore the consequences of different selective models for non-synonymous mutations. If all non-synonymous mutations are neutral, then as expected *Z* = 1 (figure 1a), and if we make some of the non-synonymous mutations deleterious, drawing their selection coefficients from a gamma distribution, as estimated from human polymorphism data (Eyre-Walker et al, 2006) we find that *Z* < 1 (figure 1a). Again, this is expected because slightly deleterious mutations (SDMs) are likely to contribute more to the level of private than shared polymorphism. If we simulate a locus in which most non-synonymous mutations are deleterious, drawn from a gamma distribution, but each locus contains a single balanced polymorphism that is shared between populations then *Z > 1* (figure 1a). It is important to note that the density of balanced polymorphisms is high in these simulations because we have simulated a short exon, of just 288bp, the average length in humans, and each one contains a balanced polymorphism. If we were to reduce the density of balanced polymorphisms then Z can be less than one even if there is balancing selection operating.

**Figure 1:**
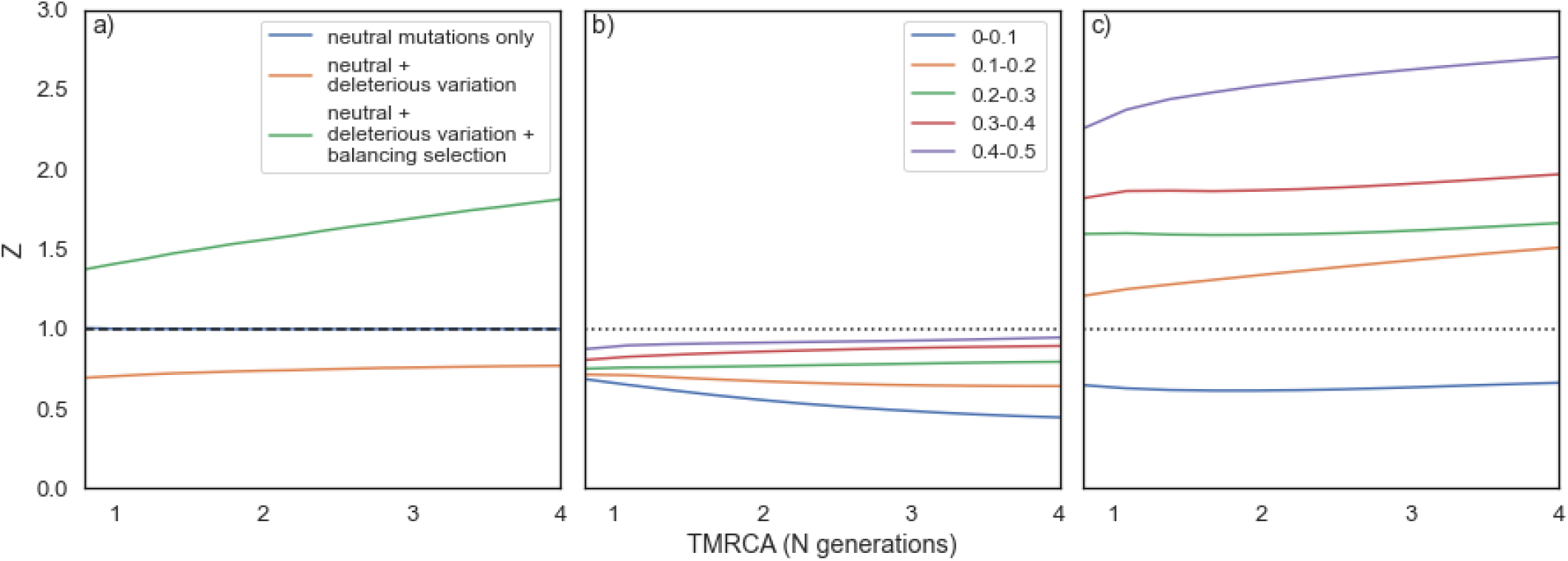
Stationary population size simulations, in which the ancestral population is duplicated to form two daughter populations of the same size to each other and the ancestor. Each simulation was repeated 2 million times. The time to the most recent common ancestor (tMRCA) is measured in N generations, where N is the population size. A Z value of greater than 1 indicates a greater proportion of shared non-synonymous polymorphisms than private non-synonymous polymorphism, which is a signal of balancing selection. For b-c) private polymorphisms have been binned by minor allele frequency, in bins of size 0.1. a) - simulations of neutral genetic variation only; a) - orange & b) neutral and deleterious variation; a) - green & c) neutral, deleterious and balanced polymorphisms.

Slightly deleterious mutations tend to depress the value of Z because they are more likely to segregate within a population, than to be shared between populations that diverged sometime in the past. There are two potential strategies for coping with this tendency. We can test for the presence of balancing selection as a function of the frequencies of the polymorphisms in the population, because SDMs will tend to be enriched amongst the rarer polymorphisms in the population. A similar approach has been used successfully to ameliorate the effects of SDMs in the classic MK approach for estimating the rate of adaptive evolution between species (Fay et al. 2001; Charlesworth and Eyre-Walker, 2008; Messer and Petrov, 2013). Or we can explicitly model the generation of shared and private polymorphisms under a realistic demographic and selection model to control for the effects of SDMs. We focus our attention here on the first of these strategies, although we touch on the latter strategy in the discussion. We apply the frequency filter to both the private and shared polymorphisms; this is necessary because if we applied the filter only to the private polymorphisms, we could be comparing high frequency private polymorphisms, with a low ratio of R_N_ to R_S_, because SDMs have been excluded, to low frequency shared polymorphisms, which may contain many SDMs and hence have a high value of S_N_/S_S_; this can yield artefactual evidence of of balancing selection. This could be exacerbated if some of the SDMs are recessive. To investigate the effects of polymorphism frequency on our estimate of Z we divided polymorphisms into 5 bins of 0.1 (we did not orient SNPs); for shared polymorphisms we estimated their frequency as the unweighted mean frequency from the two populations.

If we simulate a population in which non-synonymous mutations are deleterious, whose effects are drawn from a gamma distribution, we find that *Z* < 1 but this is less marked for the high frequency categories, as we expect. For the lowest frequency category Z decreases as a function of the time to most recent common ancestor, whereas for the higher frequency categories it is either unaffected or increases slightly (figure 1b). If we include a balanced polymorphism, introduced prior to the population split and subject to strong selection, into the model, which still also includes deleterious mutations, we find that *Z* > 1 for all frequency bins except the lowest one (figure 1c). In each case Z increases as a function of the time since the population split; this is to be expected because Z is related to the proportion of shared polymorphisms that are subject directly to balancing selection (see below), and as time progresses, so neutral and SDMs go to fixation or loss in one or both of the descendant populations. Note, once again that the level of balancing selection in these simulations is high because every locus contains a balanced polymorphism.

The simulation above does not take into account the demographic effects that a division in a population involves. We therefore performed more realistic simulations which involve vicariance and dispersal scenarios with and without migration between the sampled populations (Figures S1–S10). We also simulated with and without expansion after separation. We performed all simulations under two distributions of fitness effects (DFE), which were estimated from human and *Drosophila melanogaster* populations. In the vicariance scenario the ancestral population splits into two daughter populations of equal or unequal sizes. In the dispersal scenario a single daughter population is generated by duplicating part of the ancestral population, which remains the same size as it was before; we vary the daughter population size. In both cases, we explore the consequences of expansion after separation of the populations.

None of the simulated demographic scenarios is capable of generating Z values greater than 1 under either DFE - the method does not seem to generate false positives (Figures S1–S10). However, it is worth noting that a more severe difference in the size of the descendant populations results in depressed Z values in the smaller of the two populations, suggesting demography can affect the value of Z. In all cases the value of Z is smallest for the lowest frequency category, those polymorphisms with frequencies <0.1, and this frequency category often shows a dramatic difference to the other categories. We therefore suggest combining the polymorphisms above 0.1 when data is limited. As expected, we find that Z<1 in all simulations when we sum all polymorphisms with frequencies > 0.1 (Figures S11 and S12).

### Estimating the level of balancing selection

One of the great advantages of our method is that it gives an estimate of the number of polymorphisms that are directly affected by balancing selection, under a simple model of evolution. Let us assume that synonymous mutations are neutral and that non-synonymous mutations are strongly deleterious, neutral or subject to balancing selection; we further assume that all balanced polymorphisms arose before the two populations split. Then the expected numbers of non-synonymous, *R_N_,* and synonymous, *R_S_*, private polymorphisms are

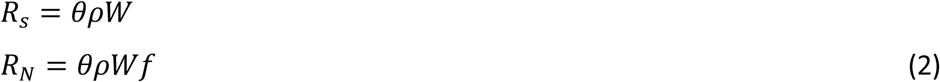

where *θ = 4N_e_u, N_e_* is the effective population size and *u* is the mutation rate per site per generation. *ρ* is the proportion of polymorphisms that are private to the population, *W* is Watterson’s coefficient and *f* is the proportion of non-synonymous mutations that are neutral, (1-*f*) being deleterious or subject to balancing selection.

In deriving expressions for *S_N_* and *S_S_* we have to take into account that a balanced polymorphism can maintain neutral variation in linkage disequilibrium that may also be shared between populations. If we have *b* balanced non-synonymous polymorphisms and each of those maintains *x* neutral mutations in linkage disequilibrium, then the expected values of *S_N_* and *S_S_* are

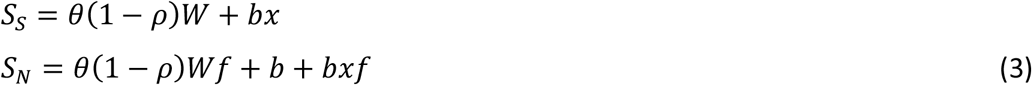

It is then straightforward to show that the proportion of shared non-synonymous polymorphisms that are directly maintained by balancing selection is

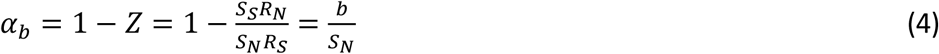

This is clearly an unrealistic model in several respects. First, it can be expected that there are slightly deleterious mutations in many populations and this will lead to an underestimation of *α_b_*; and second, it is likely that new balanced polymorphisms will be arising all the time and these will contribute to private polymorphism, increasing *R_N_/R_S_* and leading to a conservative estimate of *α_b_*.

### Data analysis - humans

We have shown that the method has the potential to detect balancing selection under realistic evolutionary models. We therefore applied our method to human data from the 1000 genomes project (The 1000 Genomes Project Consortium, 2015) focussing on four populations - Africans, Europeans, East Asians and South Asians. We find that Z >1 when private polymorphisms from the African population are used for all population comparisons if the frequency of private and shared polymorphisms >0.1; we also find that Z>1 in the South Asian and East Asian population comparison when using South Asian private polymorphisms and polymorphisms with frequencies > 0.1 (Figure 2). For several comparisons we have no polymorphisms at the relevant frequencies, and many of the confidence intervals on our estimates of Z are large. As a consequence, we combined the data for all polymorphisms with frequencies > 0.1 (Figure 2, right-most point in each panel). The patterns above are replicated; Z is significantly greater than one when we use private polymorphisms from Africa, and in the comparison between the East and South Asian populations, but Z<1 otherwise. In some cases, the Z value for the combined data can appear inconsistent with the Z values from the individual frequency categories - for example, in the European-South Asian comparison the combined Z value is greater than the Z value for 0.1-0.2 despite the fact that this is the only Z value above 0.1. This is because there are polymorphisms with frequencies >0.2, but there are not enough to yield a valid estimate of Z.

**Figure 2:**
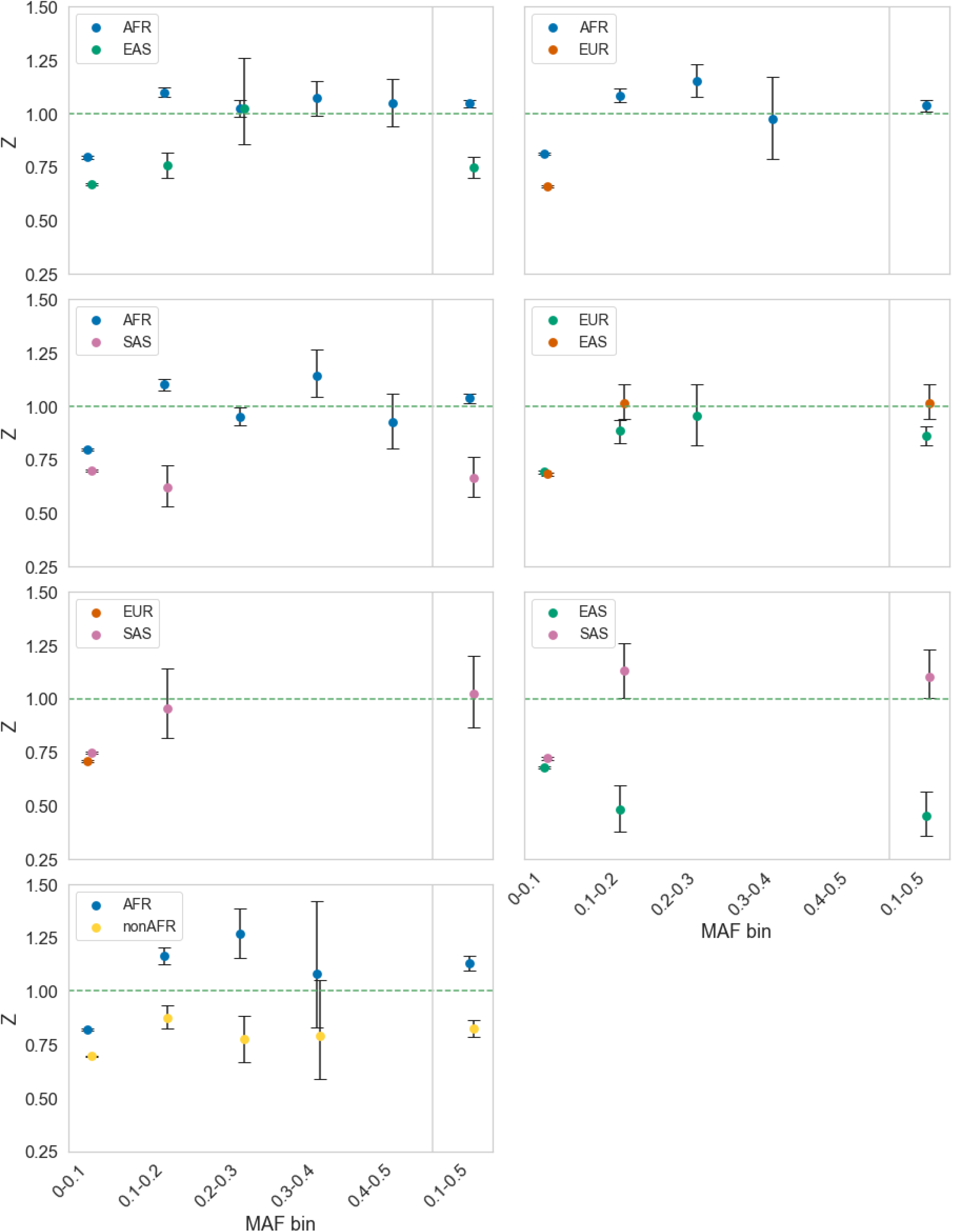
The value of Z plotted against the frequency of shared and private polymorphisms, calculated for 1000 genome data from Africans (AFR), East Asians (EAS), Europeans (EUR) and South Asians (SAS). In each panel we show the value of Z for a comparison of two populations using the private polymorphisms from each, the population used being indicated in the plot legend. Data binned by minor allele frequency bins of size 0.1 on the x-axis. Final bin is 0.1-0.5 (i.e. all data minus the lowest frequency bin). Confidence intervals were generated by bootstrapping the data by gene 100 times. Only datapoints in which there were at least 20 polymorphisms for all polymorphism categories were plotted, because the confidence intervals were very large otherwise.

Although, the values of Z are not consistently > 1 across populations, the results suggest that there is balancing selection operating. Our simulations show that Z is consistently <1 when there is no balancing selection, and that the value of Z differs between the two population comparisons if the populations have undergone different demographies. We therefore infer that there is balancing selection maintaining polymorphisms between populations.

If we estimate *α_b_* to those comparisons in which Z is significantly greater than 1, we estimate that approximately 12% of the non-synonymous shared polymorphisms between the African and other human populations are subject to balancing selection, as well as between the Asian populations (Table 1). These estimates are likely to be underestimates because there will still be SDMs segregating in our data, even though we have removed the lowest frequency variants (see simulation results). The proportions suggest that ~500 polymorphisms, which are shared between the African and other populations, are maintained by balancing selection. We estimate rather more are maintained between the two Asian populations, although the confidence intervals on this estimate are large. If we combine data across the non-African populations, we estimate that ~1400 polymorphisms are shared between African and non-African populations in total because of balancing selection (Table 1). The fact that the estimate from the African-non-African comparison is larger than the estimate from the African versus each individual population, suggests that many of the balanced polymorphisms shared between populations are unique to each population pair - i.e. there are balanced polymorphisms shared between African and European populations, that are not shared between Africans and East Asians.

**Table 1:** Estimates of the proportion of shared non-synonymous polymorphisms under balancing selection, α_b_, and the number of polymorphisms, b, being directly maintained by balancing selection for population comparisons in which Z>1. 95% confidence intervals were generated by bootstrapping the data by gene 100 times.

| Target population | Comparative population | $\alpha_b$ | $\alpha_{b\_low}$ | $\alpha_{b\_high}$ | **b** | **b<sub>low</sub>** | **b<sub>high</sub>** |
| --- | --- | --- | --- | --- | --- | --- | --- |
| African | nonAfrican | 0.12 | 0.09 | 0.14 | 1420 | 1077 | 1737 |
| African | European | 0.04 | 0.01 | 0.06 | 577 | 176 | 926 |
| African | East Asian | 0.05 | 0.03 | 0.06 | 657 | 401 | 882 |
| African | South Asian | 0.03 | 0.01 | 0.05 | 507 | 182 | 796 |
| South Asian | East Asian | 0.09 | 0.00 | 0.19 | 1273 | 18 | 2594 |

A concern in any analysis of human population genetic data is the influence of biased gene conversion (BGC). This process tends to increase the number and allele frequencies of AT>GC mutations, and reduce the number and allele frequencies of GC>AT mutations. If this process differentially affects synonymous and non-synonymous sites and shared and private polymorphisms, then it could potentially lead to Z>1. To investigate whether BGC has an effect we performed two analyses. In the first, we divided our genes according to whether they were in high and low recombining regions, dividing the data at the median recombination rate (RR). Our two groups differ substantially in their mean rate of recombination (mean RR in low group = 1.2 x 10^-7^ centimorgans per site and high group = 1.8 x 10^-6^ centimorgans per site). We find that Z is actually higher in the low RR regions, although not significantly so (Table 2), which suggests that BGC is not responsible for the comparisons in which Z > 1.

**Table 2:** Estimates of *Z* for data split by median recombination rate. Confidence intervals were generated by bootstrapping genes 100 times.

| mean recombination rate | sN | sS | rN | rS | Z | Z<sub>low</sub> | Z<sub>high</sub> |
| --- | --- | --- | --- | --- | --- | --- | --- |
| 1.20E-09 | 7023 | 7447 | 553 | 653 | 1.114 | 1.066 | 1.160 |
| 1.80E-08 | 7767 | 8514 | 604 | 702 | 1.060 | 1.021 | 1.103 |

**Table 3:**
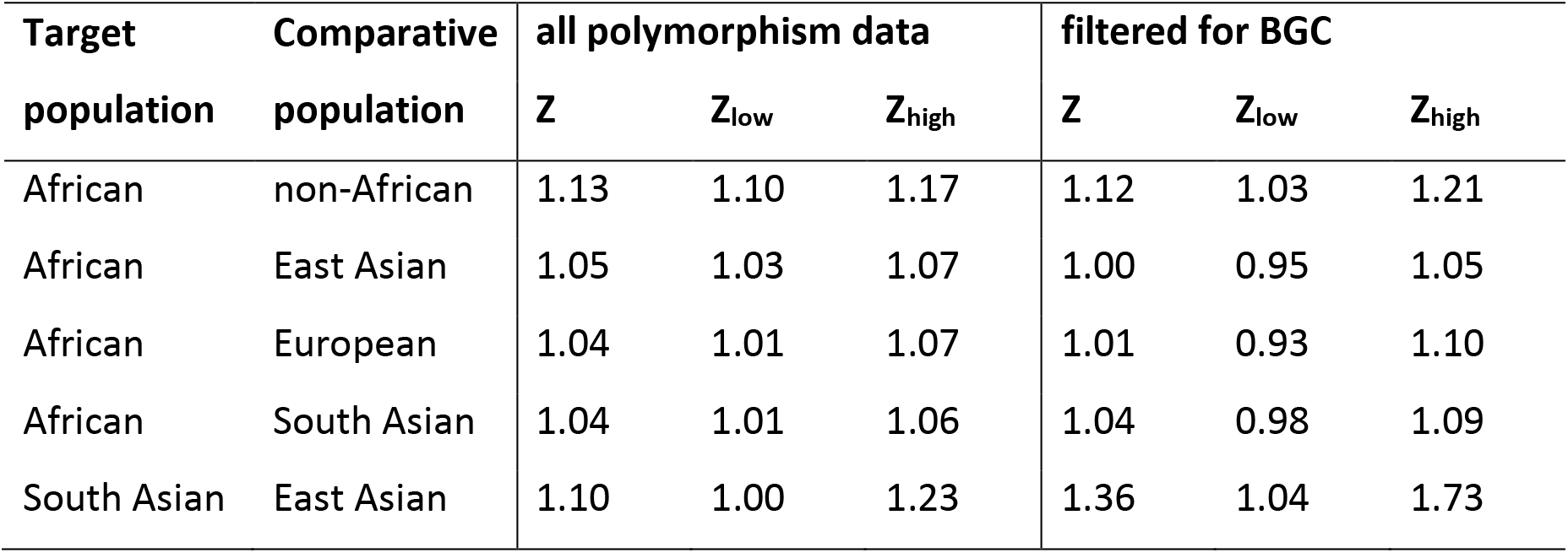
Testing the effects of BGC for population comparisons which show Z>1 using all polymorphisms with frequencies > 0.1. To control for BGC the analysis was restricted to A<>T and G<>C SNPs. 95% confidence intervals were generated by bootstrapping the data by gene 100 times.

In the second test of the influence of BGC on the value of Z we limited our analysis to mutations that are not affected by BGC - i.e. G<>C and A<>T mutations. This reduces our dataset by about 80%. As a consequence, we summed the data for all polymorphisms with frequencies >0.1. We find that our estimate is largely unchanged from that when all polymorphisms are included, however the confidence intervals are increased substantially so that Z is only significantly greater than one for the African-non-African, and the South versus East Asian comparisons (Table 2). Our two tests suggest that our results are not affected by BGC.

### Groups of genes

We can potentially apply our test of balancing selection to individual genes or groups of genes, where we have enough data. Balancing selection has been implicated in the evolution of immune related genes (e.g. Bitarello et al. 2018, Weedel and Conway, 2010, Hughes and Nei, 1988, Hedrick and Thompson, 2002), particularly major histocompatibility complex (MHC), or human leukocyte antigen genes (HLA) (Aguilar et al. 2004 and Paterson, 1998). To investigate whether we could detect th<u>is</u> signature in our data, we split our dataset into HLA and non-HLA genes **(The MHC sequencing consortium, 1999)**. Due to a lack of private polymorphisms, we combined all frequency categories >0.1. We found Z was significantly greater in HLA than non-HLA genes for all population comparisons except Europeans and South Asians using European private polymorphisms (Figure S13). However, the value of Z is greater than one in population comparisons in which African private polymorphisms are used, consistent with balancing selection maintaining variation in the HLA region. We estimate that a very substantial proportion of non-synonymous genetic variation is being maintained by balancing selection, although the confidence intervals on our estimates are large; roughly 50% of the shared non-synonymous SNPs are being maintained by balancing selection between African and non-African populations in the HLA region and this equates to approximately 200 polymorphisms (Table 4).

**Table 4:** Estimates of the proportion of shared non-synonymous polymorphisms under balancing selection, α_b_, and the number of polymorphisms being directly maintained by balancing selection, b, for population comparisons in the HLA region for population comparisons in which Z>1 when using all genes. Estimates for polymorphisms with frequency > 0.1. Confidence intervals were generated by bootstrapping the data by gene 100 times.

| target | comparative | $\alpha$ | $\alpha_{b\_low}$ | $\alpha_{b\_high}$ | $b$ | $b_{low}$ | $b_{high}$ |
| --- | --- | --- | --- | --- | --- | --- | --- |
| AFR | nonAFR | 0.702 | -0.011 | 0.779 | 299 | Na | 332 |
| AFR | EAS | 0.287 | -0.005 | 0.502 | 134 | Na | 234 |
| AFR | EUR | 0.731 | 0.317 | 0.820 | 338 | 147 | 379 |
| AFR | SAS | 0.529 | 0.307 | 0.694 | 247 | 143 | 324 |
| EAS | SAS | -0.307 | Na | 0.292 | Na | Na | 133 |

However, the signature of balancing selection that we have detected across all genes is not simply due to the HLA genes. We find that Z>1 in non-HLA genes in most population comparisons in which Z>1 for all genes, except in the comparison of East and South Asian populations (Table 5); in most cases these estimates of Z are significantly greater than 1. We estimate that >1000 of non-synonymous polymorphisms are subject to balancing selection amongst the polymorphisms shared by African and non-African populations, with several 100 shared between each of the populations and the African population.

**Table 5.**
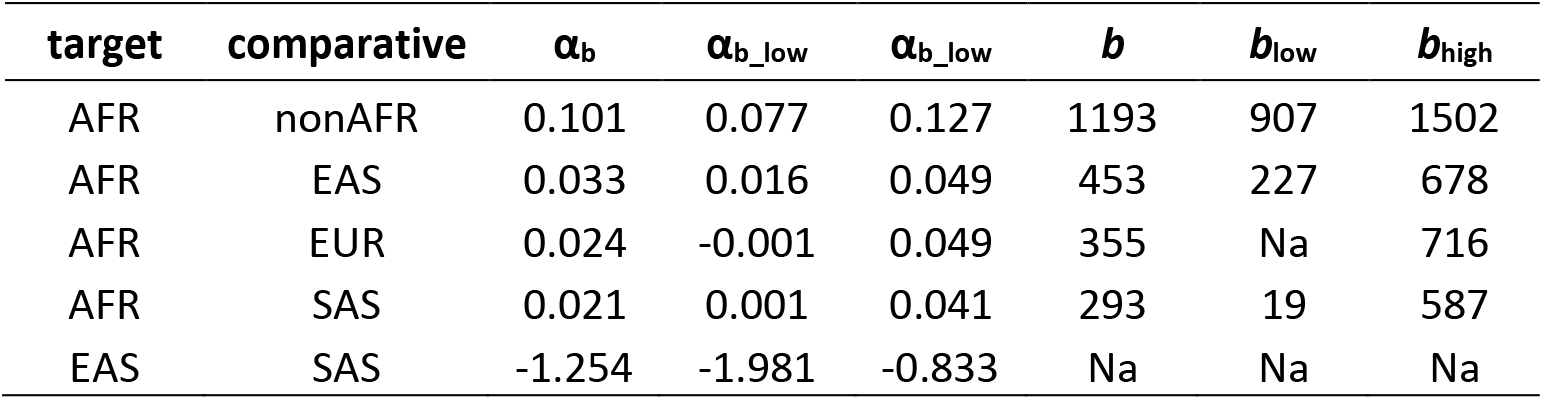
Estimates of the proportion of shared non-synonymous polymorphisms under balancing selection, α_b_, in non-HLA genes, and the number of polymorphisms being directly maintained by balancing selection, b, for population comparisons in which Z>1 when using all genes. Confidence intervals were generated by bootstrapping the data by gene 100 times.

| target | comparative | $\alpha_b$ | $\alpha_{b\_low}$ | $\alpha_{b\_high}$ | $b$ | $b_{low}$ | $b_{high}$ |
| --- | --- | --- | --- | --- | --- | --- | --- |
| AFR | nonAFR | 0.101 | 0.077 | 0.127 | 1193 | 907 | 1502 |
| AFR | EAS | 0.033 | 0.016 | 0.049 | 453 | 227 | 678 |
| AFR | EUR | 0.024 | -0.001 | 0.049 | 355 | Na | 716 |
| AFR | SAS | 0.021 | 0.001 | 0.041 | 293 | 19 | 587 |
| EAS | SAS | -1.254 | -1.981 | -0.833 | Na | Na | Na |

If we run our analysis grouping genes by their GO category and restricting the analysis to those groups that have at least 100 polymorphisms with frequencies >0.1 we find 683 categories in which Z is significantly greater than 1 in at least population comparison (Supplementary file S1) and we list those significant in 5 or more population comparisons in Table 6. One of these GO categories, “nucleic acid binding” is shared across 7 of the 14 population comparisons, “endoplasmic reticulum membrane” across 6 population comparisons; amongst those categories shared among 5 are “viral process” and “immune system process”, but there are many others which are surprising including “protein import into the nucleus” (Supplementary file S1). Eighty-five categories are shared between 4 or more population comparisons, and 155 amongst three or more population comparisons. These include 7 categories related to immunity (including immune system process which is significant in 5 population comparisons), and 40 categories that are linked to antigen presentation though not classified as immune-related categories. There are also 5 viral-related categories (including viral process which is significant in 5 population comparisons).

**Table 6.**
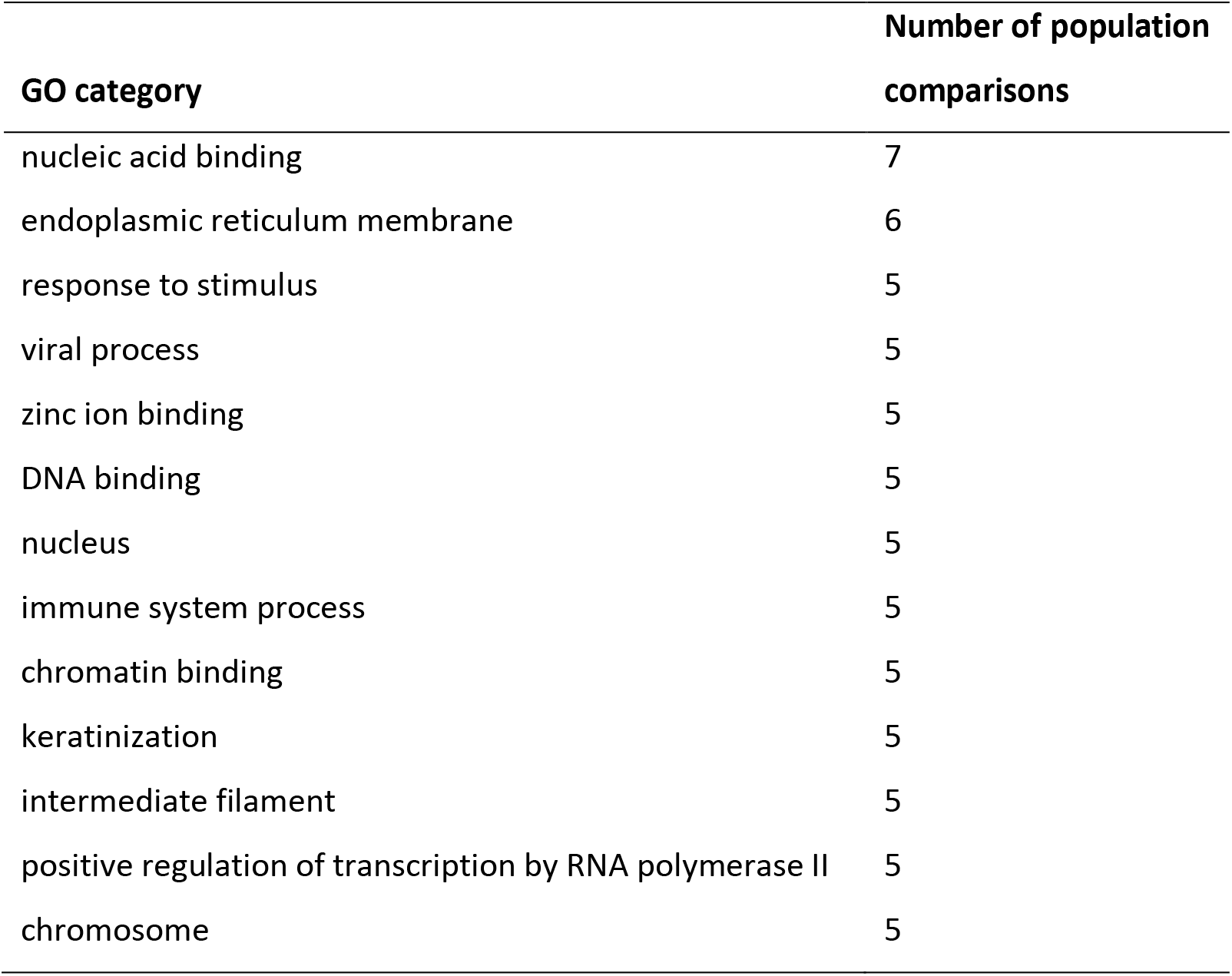
GO categories in which Z is significantly greater than one in at least 5 population comparisons.

### Individual genes

We applied our statistic to individual human genes, combining all frequency bins (0-0.5) due to a lack of polymorphism. We tested for significance using a one-tailed Fisher’s exact test. Of the 14,261 genes we analysed 514 had Z>1 in at least one population comparison. Eighteen of these were nominally significant at p<0.1 (supplementary file S2), but no gene was individually significant when we corrected for multiple testing using a Bonferroni correction. Eighteen genes have Z>1 in at least 9 population comparisons; note that since populations share polymorphisms, we cannot combine the evidence for balancing selection across these populations (Table 7). Four of these genes MUC4, RP1L1, PKD1L2 and ZAN have Z>1 in all population comparisons.

**Table 7.** Genes with Z>1 in multiple population comparisons.

| Gene symbol | Number of population comparisons in which $Z > 1$ |
| --- | --- |
| MUC4 | 14 |
| RP1L1 | 14 |
| PKD1L2 | 14 |
| ZAN | 14 |
| C1orf167 | 13 |
| SPTBN5 | 12 |
| MKI67 | 12 |
| DNAH14 | 11 |
| WDFY4 | 10 |
| FAM230G | 10 |
| CMYA5 | 9 |
| CRIPAK | 9 |
| SYNE2 | 9 |
| FSIP2 | 9 |
| GREB1 | 9 |
| ALMS1 | 9 |
| MUC19 | 9 |
| CENPF | 9 |

If we use the 514 genes and do a GO enrichment analysis, we find multiple GO categories enriched for these genes including immune response categories with 3-fold enrichment. The most highly enriched categories are involved in energy production and conversion (including dynein binding) and intracellular transport (including microtubule motor activity) (Supplementary file S2).

## Discussion

We propose a new method for detecting and quantifying the amount of balancing selection that is operating on polymorphisms, in which the numbers of non-synonymous and synonymous polymorphisms that are shared between populations and species are compared to those that are private. The method is analogous to the McDonald-Kreitman test used to test and quantify the amount of adaptive evolution between species (McDonald & Kreitman, 1991). We show that our test is robust to the presence of slightly deleterious mutations under simple demographic models of population division, expansion and migration. When we apply our method to data from human populations, we find evidence that hundreds of non-synonymous polymorphisms are being directly maintained by balancing selection in human populations.

Our method for detecting balancing selection is simple to apply and appears to be robust to changes in demography. The classic MK test of adaptive evolution between species can generate artefactual evidence of adaptive evolution if there are SDMs and there has been population size expansion (McDonald & Kreitman, 1991; Eyre-Walker, 2002); this is because SDMs that might have been fixed when the effective population size was small, no longer segregate once the population size is large. A similar bias does not appear to affect our test, although we have only investigated two DFEs and a limited number of demographic scenarios. Our test is likely to be more robust than the classic MK test because the shared polymorphisms are affected by the demographic changes that affect the private polymorphisms; i.e. if the population expands this will increase the effectiveness of natural selection on both the private and the shared polymorphisms. However, although our method seems to be relatively robust to changes in demography, in the sense that it does not generate artefactual evidence of balancing selection, it is evident that demography does affect the chance of balancing selection being identified, because the values of Z depend on the demography and which population the private polymorphisms are taken from (Figure 2).

The method can in principle be applied to any pair of populations or species. However, the test is likely to be weak when the populations/species are closely related for two reasons. First, there will be relatively few private polymorphisms, and second, the proportion of shared polymorphisms that are subject to balancing selection is likely to be low, because so many neutral polymorphisms are shared between populations because of recent common ancestry. As the populations/species diverge so the number of private polymorphisms will increase, and the proportion of shared polymorphisms that are balanced will increase. Of course, as the time of divergence increases so the selective conditions that maintained the polymorphism are likely to change and the polymorphism might become neutral, or subject to directional selection. Our method is also likely, like all methods, to be better at detecting balanced polymorphisms that are common, because most populations are dominated by large numbers of rare neutral variants. Finally, our method requires that the neutral and selected sites are interdigitated; the method is therefore easy to apply to protein coding sequences, but may be more difficult to apply to other types of variation, such as that which affects gene expression.

A great advantage of our method is that it gives an estimate of the proportion and number of shared polymorphisms that are directly subject to balancing selection, under a set of simplifying assumptions. However, the method is likely to yield underestimates of the proportion of balanced polymorphisms, under a more realistic models of evolution. We have assumed, in deriving α_b_, that all non-synonymous mutations are either strongly deleterious, neutral or subject to balancing selection. However, a substantial fraction of non-synonymous mutations appear to be slightly deleterious in humans (Cargill et al, 1999; Fay et al, 2001; Eyre-Walker et al, 2002; Hughes, et al, 2003; Asthana, et al., 2007) and other species (Fay et al, 2001; Eyre-Walker et al, 2002; Hughes, 2005; Charlesworth & EyreWalker, 2006) - i.e. they are deleterious, but sufficiently weakly selected that they contribute to polymorphism. Under stationary population size assumptions - i.e. in which the ancestral population is duplicated to form the daughter populations - this will lead to an underestimate of Z because SDMs tend to contribute more to private than shared polymorphism, and hence inflate R_N_/R_S_ relative to S_N_/S_S_ (Figure 1). Under more realistic demographic models, in which at least one of the derived populations is reduced, this is expected to depress Z in the population that is being reduced because more SDMs will tend to segregate in smaller populations hence inflating R_N_/R_S_ (compare Figure 1 to Figures S2 and S3). Simulations suggest there is however a slight increase in Z using private polymorphisms from the larger of the populations but these Z values never exceed 1 (Figures S2 and S3). The second reason that we are likely underestimating the number of balanced polymorphisms using our simple method is that we assume that there are no balanced polymorphisms that are private to each population; these would inflate R_N_/R_S_. Private balanced polymorphisms might arise from an ancestral polymorphism that is lost from one of the daughter populations, or one that arises de novo. A more realistic model balancing selection is one in which balanced polymorphisms are continually generated with the selective forces persisting for some time before they dissipate (Sellis et al, 2011) and the balanced polymorphism is lost. The process of population division itself is likely to lead to the loss of many balanced polymorphisms as the environment shifts in the two daughter populations.

A potential solution to the tendency for our method to underestimate Z is to simulate data under a realistic demographic model assuming that there is no balancing selection, and interpret the observed values of Z that are greater than simulated values, as evidence of balancing selection. A similar approach has been used to estimate the rate of adaptive evolution between species (e.g. Eyre-Walker & Keightley, 2009; Boyko, et al., 2008; Schneider et al, 2011; Galtier, 2016; Tataru et al, 2017). However, there are challenges in this approach; in particular, we need an accurate demographic model. We have performed simulations under the commonly used human demographic model inferred by Gravel et al. (2011) estimating the DFE from the current African population; we chose the African population because it has been subject to relatively modest demographic change. Our observed Z values do not match the simulated values (Figure S14); in particular we find that the observed values of Z are substantially greater than the simulated amongst the low frequency polymorphisms. However, the model of Gravel et al. does not fit the SFS of the individual populations of 1000 genome data; for example, in the African population there are far too many singleton SNPs even amongst the putative neutral synonymous mutations (Figure S15). The lack of fit is perhaps not surprising; Gravel et al. inferred their model using 80 chromosomes per population, whereas the 1000 genome data contains >1000 chromosomes per population. Furthermore, the inference of a demographic model should take into account the influence of biased gene conversion and background selection, which appear to be pervasive factors in the human genome (Pouyet et al, 2018), so these simulations will be complex.

It might be argued that the evidence of balancing selection is weak because we typically find Z>1 using the private polymorphism from only one of the two populations. However, we have been unable to find a demographic model in which there is no balancing selection and Z>1 - note that we never observe Z>1 under the Gravel et al. model of human demography even when we change the parameters of the DFE and demographic model; furthermore, we find that simulations which involve different demographies in the two populations generate different Z values for the two populations, so there is an expectation in many species that we will observe Z values that differ between populations.

Values of Z in excess of one could potentially be due to biased gene conversion; we expect BGC to increase the allele frequency of AT>GC mutations and to decrease the frequency of GC>AT mutations; this will tend to make AT>GC mutations more likely to be shared between populations than GC>AT mutations. Since the GC-content at the third codon position is typically higher than the GC content at the first two positions, this will mean that there are more non-synonymous AT>GC mutations than synonymous, and hence more shared non-synonymous than synonymous polymorphisms. However, our results do not appear to be affected by BGC; results are similar between genes in high and low recombination rate regions of the human genome (Table 2), and our point estimates of Z are largely unaffected by restricting the analysis to SNPs which are unaffected by BGC, although the confidence intervals increase substantially (Table 3).

We estimate that there ~500 balanced polymorphisms shared between the African and each of the other human populations, ~1250 shared between the Asian populations and ~1400 shared between the African and non-African populations; these are likely to be underestimates due to the presence of slightly deleterious mutations. The fact that we estimate that ~1400 shared non-synonymous polymorphisms are being maintained between African and non-African populations, but only ~500 between each of the individual populations and the African population suggests, that many of the polymorphisms shared between the African and each individual population are unique - i.e. many polymorphisms shared between Africans and Europeans, may not be shared between Africans and Asians. These numbers are substantial, but are consistent with those of Bitarello et al. (2018) who estimated that ~8% of human genes show some evidence of long-term balancing selection, and many of these signatures are shared between populations. However, their method could not determine whether the balanced polymorphisms were coding or noncoding mutations.

As expected, we find evidence of balancing selection affecting the HLA or MHC genes (table 4) (Hughes & Nei; 1988; Hedrick, 1998). However, we find evidence of balancing selection even when these genes are removed from the analysis (Table 5). The analysis of GO categories shows that numerous categories show evidence of balancing selection across multiple population comparisons (supplementary file S1). Some of these are expected, but many are not, such as “nucleic acid binding”, which is significant in 7 of the 14 population comparisons.

No individual gene is significant when we control for multiple testing, however, several genes have Z>1 in multiple population comparisons including 10 which are shared across at least 10 of the 14 population comparisons. Three of these overlap with previous genomewide scans of selection, namely the protein-coding gene DNAH14, implicated in brain compression and encoding axenomal dynein (Voight et al, 2006); MUC4, implicated in biliary tract cancer (Tennessen & Akey, 2011); and ZAN, which encodes a protein involved in sperm adhesion, previously implicated in balancing selection and positive selection in human populations (Gasper & Swanson, 2006). Two of these ten genes are associated with tumours. MKI67 expression is associated with a higher tumour grade and early disease recurrence (Yang, et al., 2017), and WDFY4 plays a critical role in the regulation of certain viral and tumour antigens in dendritic cells (Theisen, et al., 2019). PKD1L2 is associated with polycystic kidney disease and RP1L1 variants are associated with several retinal diseases including occult macular dystrophy (Davidson, et al., 2013). SPTBN5 encodes for the cytoskeletal protein spectrin, that plays a role in maintaining cytoskeletal structure (Huh, Glantz, Je, Morrow, & Kim, 2001) and C1orf167 expresses open reading frame protein that is highly expressed in the testis (Fagerberg, et al., 2014). Finally, FAM230G is highly expressed in testes (Delihas, 2018).

Twenty-five of the 514 with Z>1 genes overlap with those genes identified by Bitarello et al. (2018), but this is similar to the level of overlap expected at random; i.e. they observed that 7.9% of protein coding genes overlapped regions identified by their method as being subject to balancing selection, and we identified 514 candidates; so we expect 0.079 x 514 = 41 by chance alone. The lack of a significant overlap is possibly not surprising; we have applied our method to non-synonymous variation, whereas the method of Bitarello et al. (2018) considers all variation. Furthermore, the method of Bitarello et al. (2018) is most powerful at detecting balancing selection over long time periods; in the case of humans, over periods of millions of years. In contrast, we have applied our method to populations that diverged 10,000’s of years ago.

It is possible that the signature of balancing selection is caused by a form of associated overdominance, in which neutral alleles at a locus are linked to different deleterious recessive alleles at other loci. For example, let us imagine that we have two closely linked loci at which we have deleterious alleles; let the A2 allele be the recessive allele at the A locus and the B2 allele at the B locus. Now consider a third neutral locus with alleles C1 and C2. If C1 is in linkage disequilibrium (LD) with the A2 allele, and C2 is in LD with the B2 allele, then C1C2 heterozygous individuals will have higher fitness than C1C1 and C2C2 homozygotes. This form of selection can lead to the maintenance of genetic variation (Zhao and Charlesworth 2016) in low recombination rate regions. However, Z is not substantially greater in regions of low recombination so AOD seems an unlikely explanation (Table 2).

## Conclusion

We present a new approach to test for the presence of balancing selection and to the number of polymorphisms that are directly affected by it. Our method appears to be robust to demographic change. Application of the method to human population genetic data suggests that 100s of non-synonymous polymorphisms shared between populations are being maintained by balancing selection.

## Methods and Materials

### Human data

Human variation data was obtained from 1000 genomes Grch37 vcf files (The 1000 Genomes Project Consortium, 2015). Variants were annotated using Annovar’s hg19 database (Wang & Li, 2010). The annotated data was then parsed to remove multi-nucleotide polymorphisms and indels. Because 1000 genomes data provides allele frequencies for the non-reference allele rather than the minor allele, the minor allele frequency for each superpopulation and also for the global minor allele frequency was calculated. We used 1000 genomes from the African, South Asian, East Asian and European populations. The American population was removed due to the fact that it is an admixed population. GO category information was obtained from Ensembl’s BioMart data mining tool (Yates, et al., 2020). We used pyrho demography-aware recombination rate maps (Spence & Song, 2019) for analyses that control for recombination rate.

### Simulations

All simulations were run using the SLiM software platform (Haller & Messer, 2017). Parameter values were taken from human estimates. Almost all simulations were of a 288bp locus, this being the average size of a human exon (Yates et al. 2020). Unless otherwise stated, the scaled recombination rate and scaled mutation rate were set at *r* = 1.1 x 10^-8^ (Dumont & Payseur, 2008)^;^ *μ*= 2.5 x 10^-8^ (Nachman & Crowell, 2000)in the ancestral population. The distribution of fitness effects was assumed to be a gamma distribution and the shape and mean strength of selection estimates for humans were taken from Eyre-Walker et al. (2006) (β = 0.23; mean *N_e_s* = 425). For *Drosophila* estimates were taken from Keightley and Eyre-Walker (2007) (β = 0.35; mean *N_e_s* = 1800); again these were values in the ancestral population. Unless dominance was fixed, it was calculated using the model of Huber et al. (2018), which was estimated from Arabidopsis. The Huber model varies the dominance coefficient depending on the selection coefficient of the mutation, where the dominance coefficient increases with the strength of selection. It’s formula is 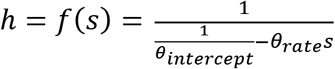 where *θ_intercept_* defines the values of *h* at *s* = 0 and *θ_rate_* determines how quickly *h* approaches zero with decreasing negative selection coefficient. We set *θ_intercept_* to 0.5 so that all mutations with a selection coefficient of *s* = 0 have a dominance coefficient, *h* = 0.5, and *θ_rate_* = 41225.56. This assumes an inverse relationship between *h* and *s*, which gives the highest log likelihood score of the relationships compared by Huber et al. (2018). For balancing selection simulations, we assume a model of heterozygote advantage and the strength of selection was sampled from a distribution such that the equilibrium frequency was uniformly distributed between 0 and 1; however, it should be noted that some balanced polymorphisms with low equilibrium frequencies were lost in one of the descendent populations. These simulations were discarded. The balanced polymorphism is introduced at the centre of the 288bp region. Two million successful simulation runs were conducted for each model.

For the generic simulations (i.e. not those involving the human demographic model) the ancestral population size was set at 200. This was allowed to equilibrate for 15N generations before a balanced polymorphism was introduced 5N generations before the population was split into two. The descendant populations were then sampled every 0.05N generations up to 20N generations after the split. We ran five different generic simulations: (i) simulations in which the ancestral population was duplicated, (ii) vicariance simulations in which the ancestral population was divided between the daughter populations in splits of 0.5N-0.5N; 0.75N-0.25N; 0.9N-0.1N, (iii) variance simulations in which the descendant populations expanded, iv) dispersal simulations, in which some variable fraction (0.5N; 0.25N; 0.1N) of the ancestral population is duplicated to form the dispersal population, and the ancestral population continues as the other daughter population, and v) dispersal with population increase of the dispersal population. The dispersal population starts as 0.1N and expands exponentially 2 to 10x its original size after 21N generations. Scenarios ii-v were repeated with migration rates of 0.01N and 0.001N of the ancestral population size between the descendant populations.

We also ran some simulations under the human demographic model of Gravel et al. (2011); for details of the demographic structure of the simulation (Gravel et al. 2011). The distribution of fitness effects for deleterious mutations was assumed to be a gamma distribution using the parameters estimated from the African superpopulation using the GammaZero model within the Grapes software (Galtier, 2016); the parameters are similar to those estimated by Eyre-Walker et al (2006), and used in the generic simulations (Gamma shape = 0.17 and Mean *N_e_S* = 1144). We chose to infer the DFE for the African superpopulation because this is currently the largest dataset available for a population that has been inferred to be relatively stable. Dominance was calculated using the Huber model discussed above. Sampling of all populations (African, East Asian and European) was conducted at the end of the simulation (i.e. the equivalent of the present day).

## Acknowledgements

the work was supported by grant #????? from the Natural Environment Research Council to MV.

## Supplementary Material

**Supplementary Table 1:**
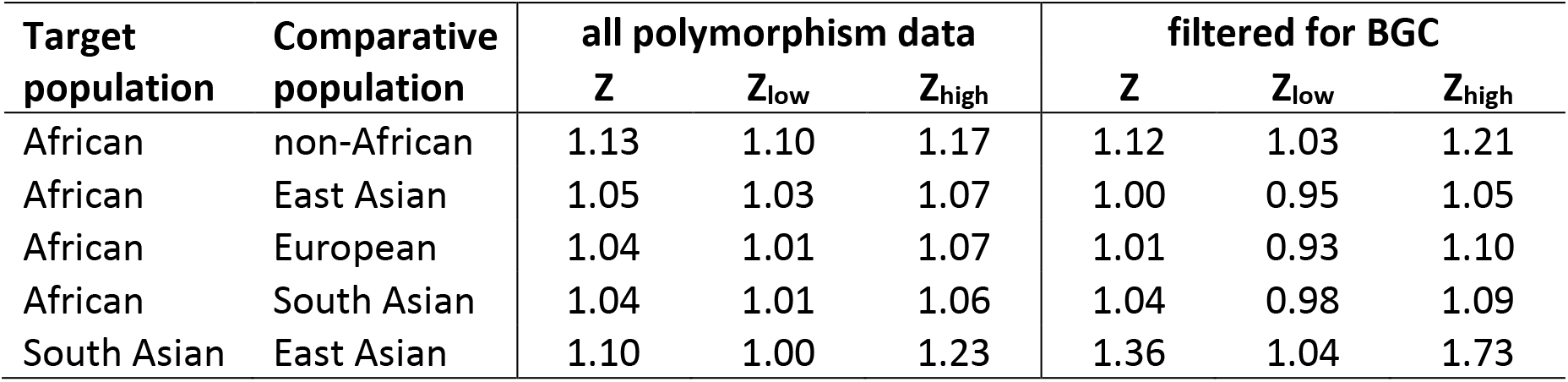
Testing the effects of BGC for population comparisons which show Z>1. Confidence intervals were generated by bootstrapping the data by gene 100 times.

**Supplementary Table 2:** Estimates of Z for HLA genes only. Confidence intervals were generated by bootstrapping the data by gene 100 times.

| target | comparative | **Z** | **Z<sub>low</sub>** | **Z<sub>high</sub>** | **$\alpha$** | **$\alpha_{b\_low}$** | **$\alpha_{b\_low}$** | ***b*** | ***b<sub>low</sub>*** | ***b<sub>high</sub>*** |
| --- | --- | --- | --- | --- | --- | --- | --- | --- | --- | --- |
| AFR | nonAFR | 3.354 | 0.989 | 4.533 | 0.702 | -0.011 | 0.779 | 299 | Na | 332 |
| AFR | EAS | 1.403 | 0.995 | 2.007 | 0.287 | -0.005 | 0.502 | 134 | Na | 234 |
| AFR | EUR | 3.714 | 1.465 | 5.568 | 0.731 | 0.317 | 0.820 | 338 | 147 | 379 |
| AFR | SAS | 2.123 | 1.443 | 3.272 | 0.529 | 0.307 | 0.694 | 247 | 143 | 324 |

**Supplementary Table 3:** Estimates of Z for non-HLA genes only. Confidence intervals were generated by bootstrapping the data by gene 100 times.

| target | comparative | **Z** | **Z<sub>low</sub>** | **Z<sub>high</sub>** | **α** | **α<sub>b_low</sub>** | **α<sub>b_low</sub>** | **b** | **b<sub>low</sub>** | **b<sub>high</sub>** |
| --- | --- | --- | --- | --- | --- | --- | --- | --- | --- | --- |
| AFR | nonAFR | 1.112 | 1.083 | 1.146 | 0.101 | 0.077 | 0.127 | 1193 | 907 | 1502 |
| AFR | EAS | 1.034 | 1.017 | 1.052 | 0.033 | 0.016 | 0.049 | 453 | 227 | 678 |
| AFR | EUR | 1.025 | 0.999 | 1.052 | 0.024 | -0.001 | 0.049 | 355 | Na | 716 |
| AFR | SAS | 1.021 | 1.001 | 1.043 | 0.021 | 0.001 | 0.041 | 293 | 19 | 587 |

**Supplementary Figure S2:**
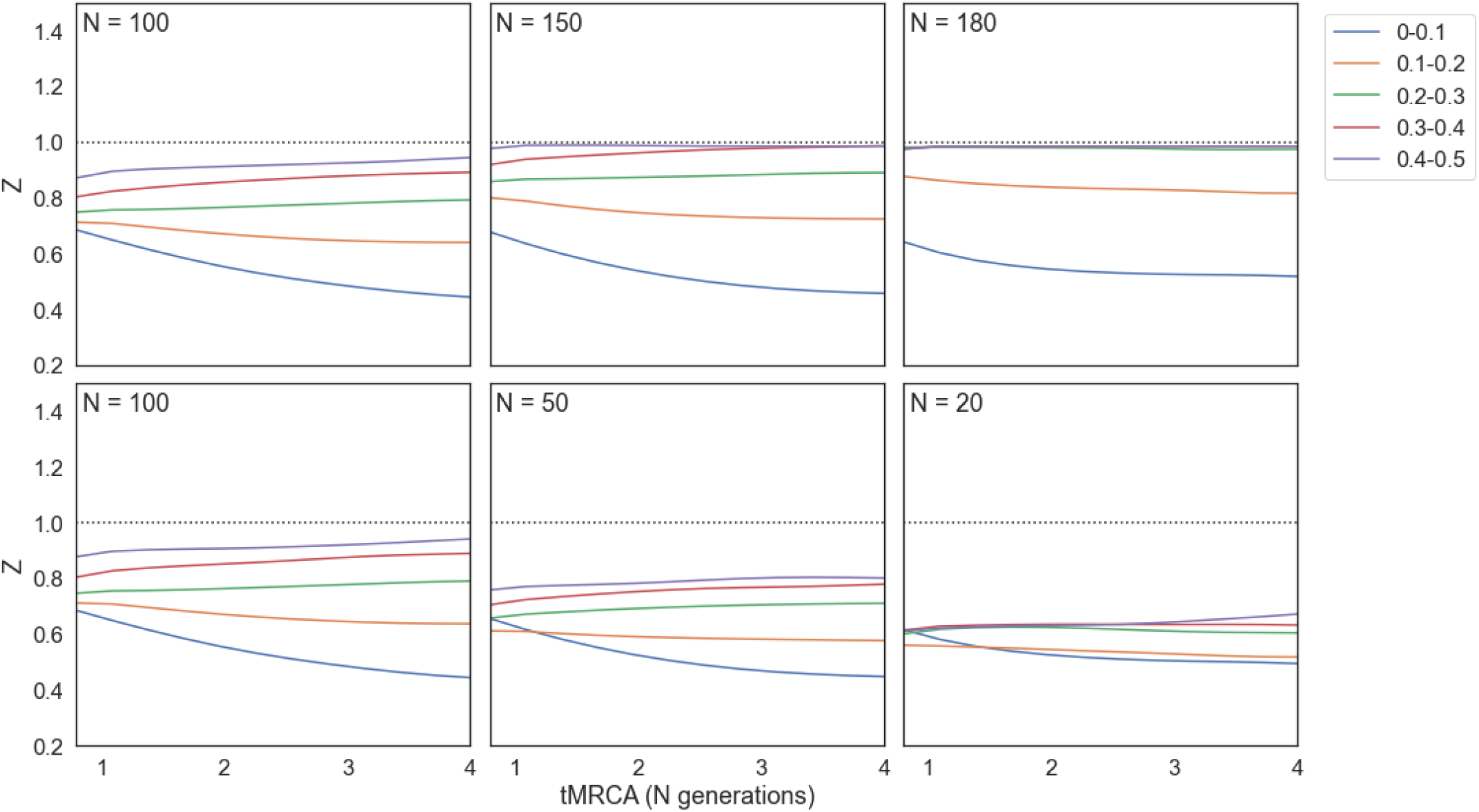
Vicariance simulations in which the ancestral population splits to form two daughter populations of the size specified in the panel. Each column is a separate set of simulations, with the top row plotting Z against tMRCA (measured in N generations, where N is the population size) for the larger daughter population, and the bottom row the smaller. Deleterious mutations are drawn from a gamma DFE with parameters inferred from human population data.

**Supplementary Figure S2:**
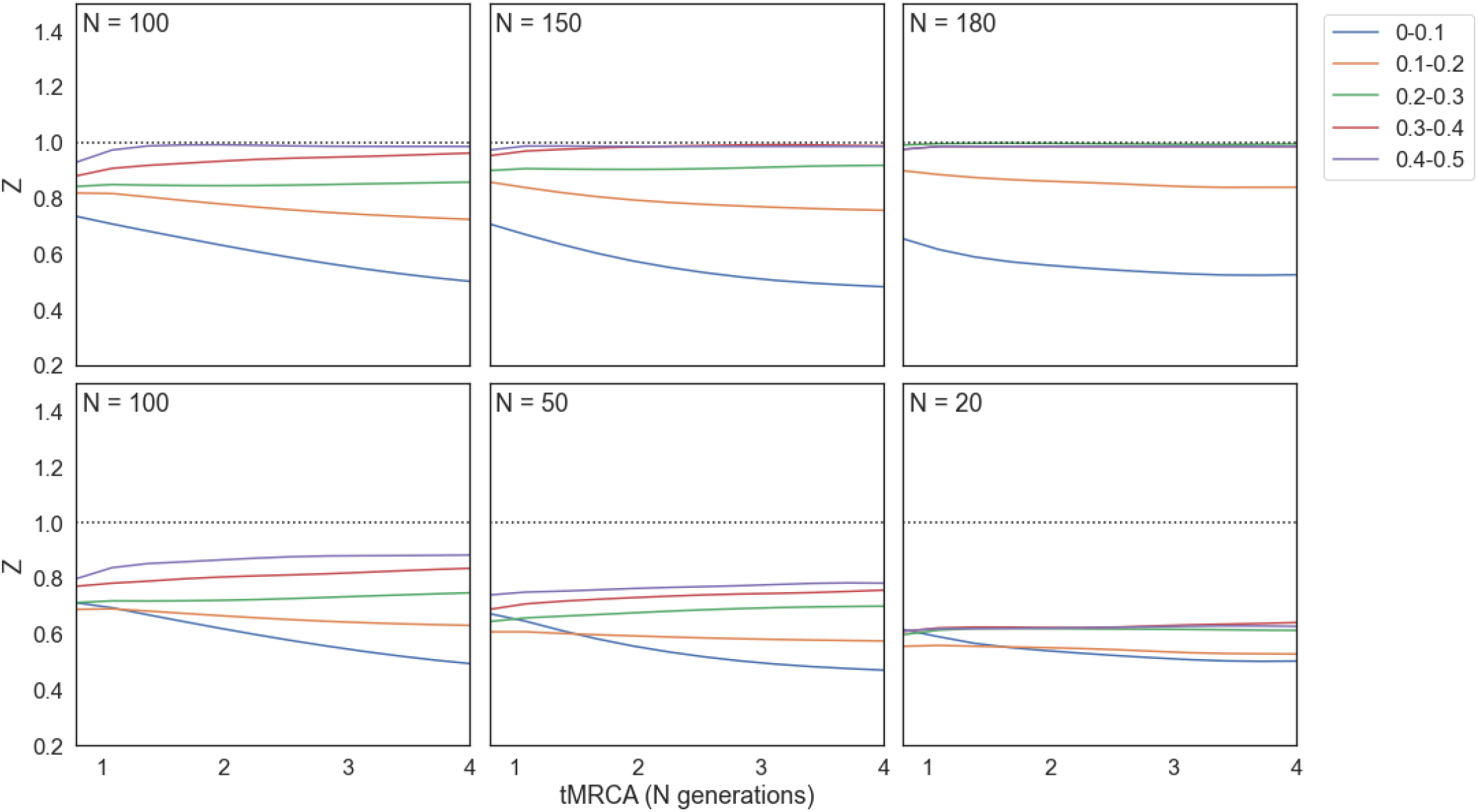
Dispersal simulations in which a single daughter population disperses from the ancestral population. Each column is a separate set of simulations, with the top row plotting Z against tMRCA (measured in N generations, where N is the population size) for the ancestral population, and the bottom row the daughter population. Deleterious mutations are drawn from a gamma DFE with parameters inferred from human population data.

**Supplementary Figure S3:**
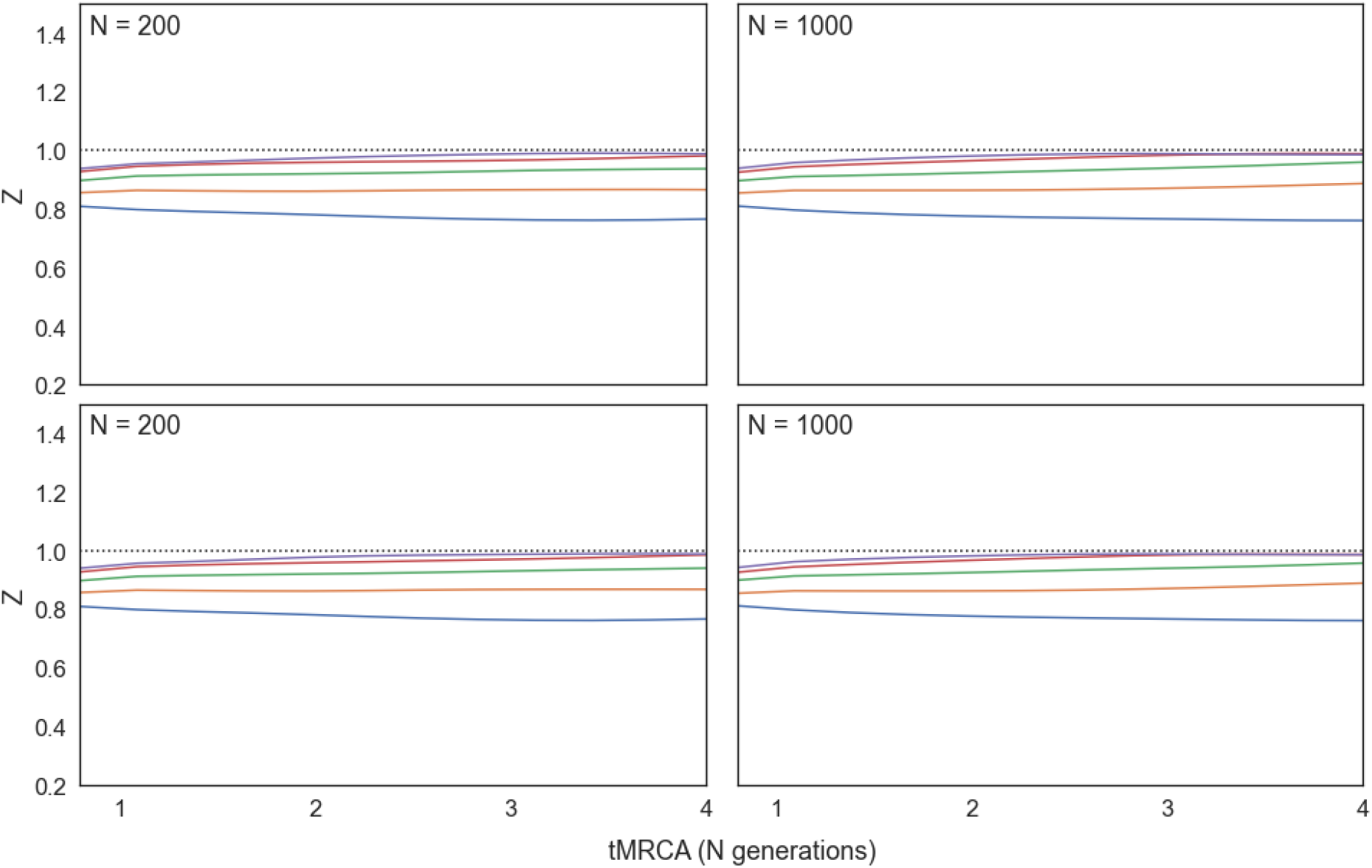
Vicariance expansion simulations in which both daughter populations expand. The ancestral population (of size N=200) splits to form two daughter populations of size N=100. Both daughter populations go on to expand in size. In the left column the daughter populations double in size. In the right panel they reach 10x their initial size. Deleterious mutations are drawn from a gamma DFE with parameters inferred from human population data.

**Supplementary Figure S4:**
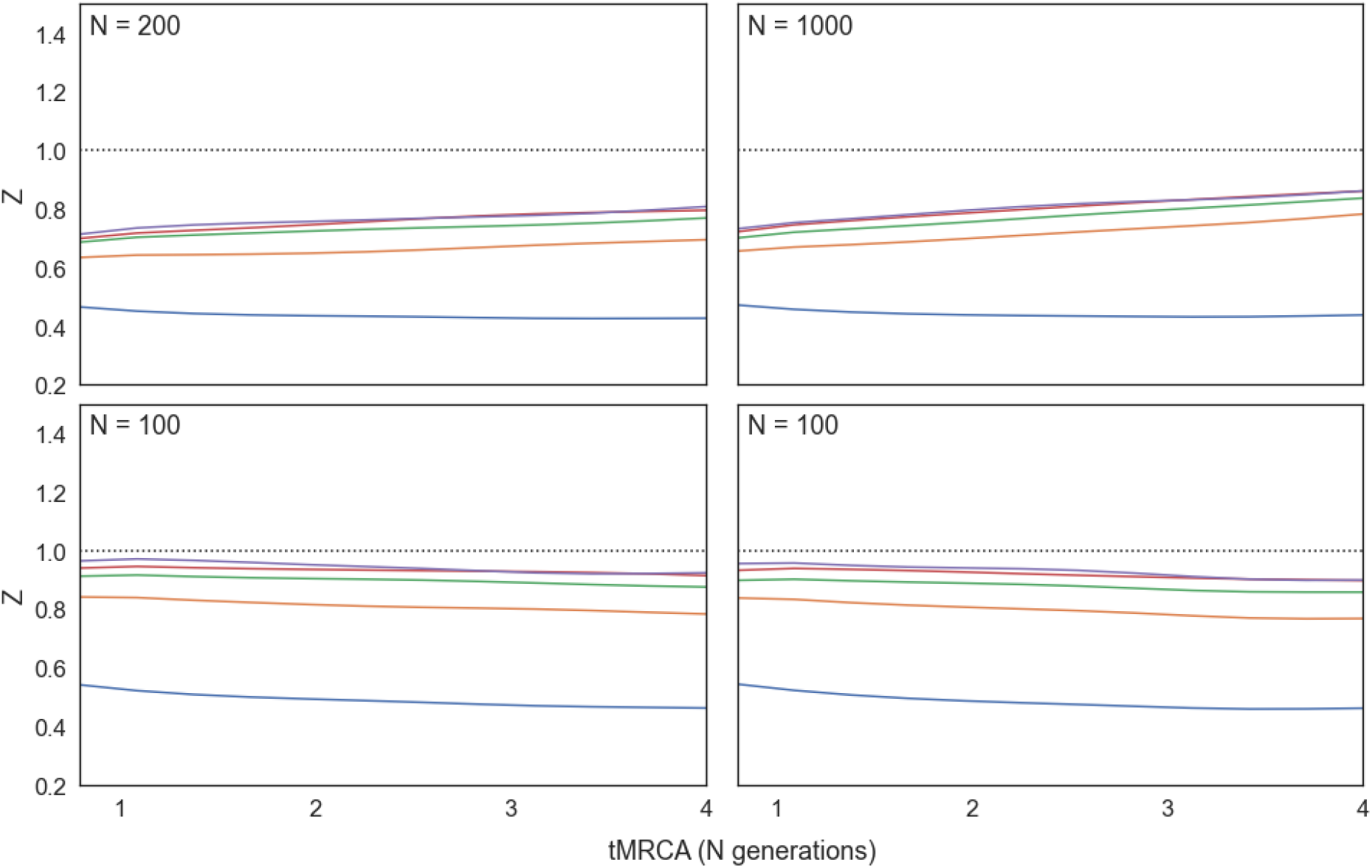
Vicariance expansion simulations in which only one daughter population expands. The ancestral population (of size N=200) splits to form two daughter populations of size N=100. One daughter population (upper panels) goes on to expand in size. In the left column the daughter populations double in size. In the right panel they reach 10x their initial size. Deleterious mutations are drawn from a gamma DFE with parameters inferred from human population data.

**Supplementary Figure S5:**
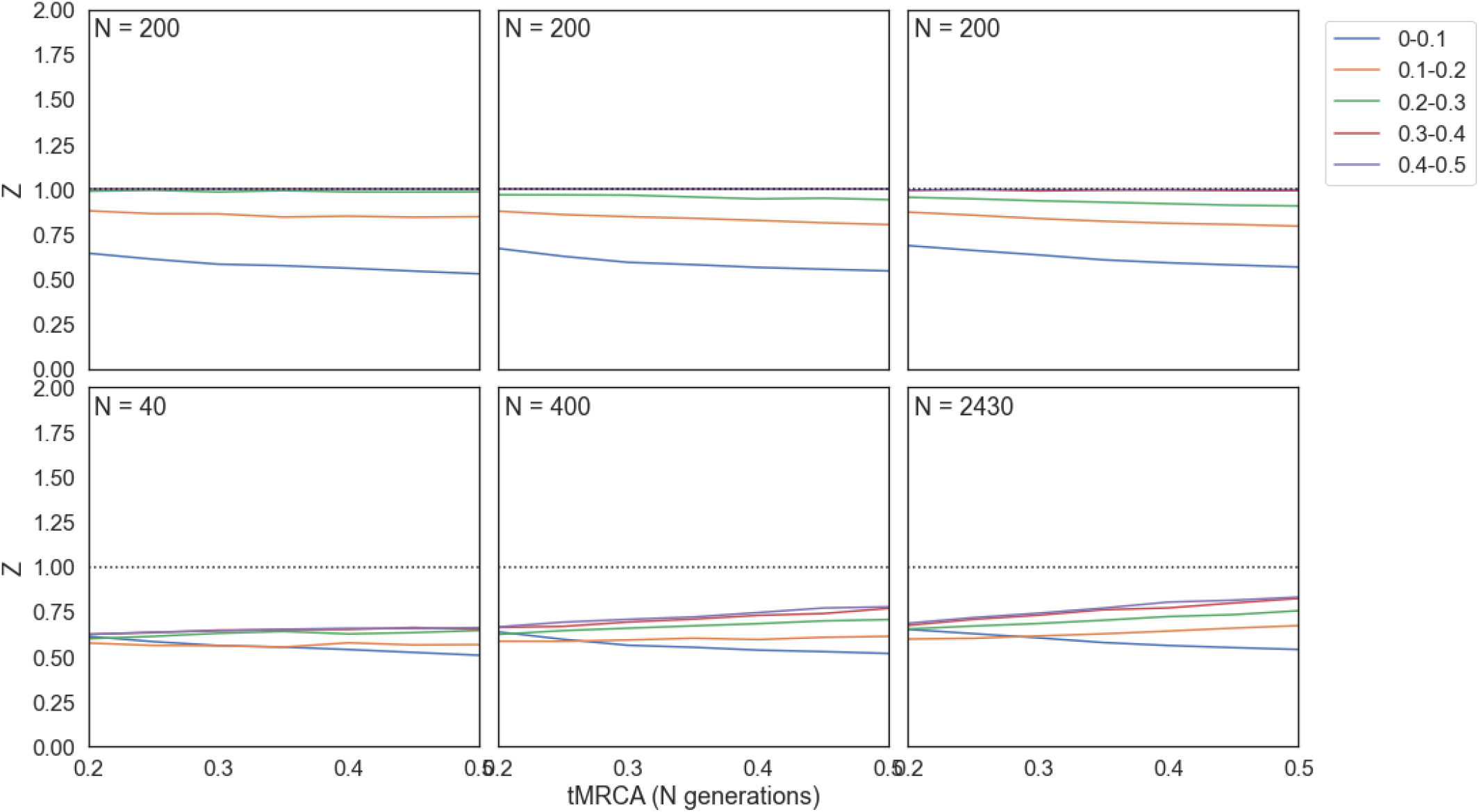
Dispersal expansion simulations in which a single daughter population disperses from the ancestral population and then expands. The ancestral population (of size N=200) splits to form a daughter population of size N=100, which expands to the final population size shown in the panel. Each column is a separate set of simulations, with the top row plotting Z against tMRCA (measured in N generations, where N is the population size) for the ancestral population, and the bottom row the daughter population. Deleterious mutations are drawn from a gamma DFE with parameters inferred from human population data.

**Supplementary Figure S6:**
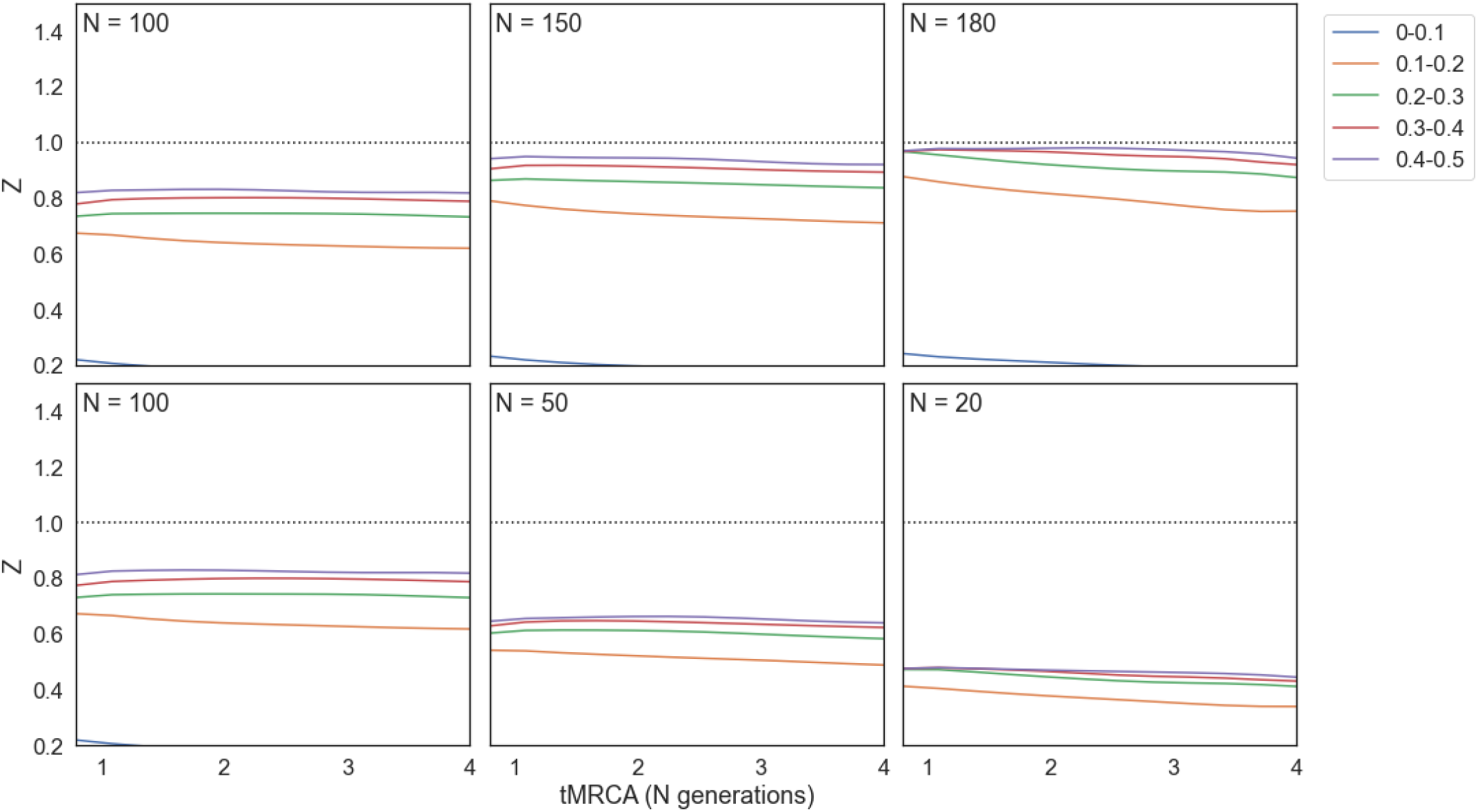
Vicariance simulations in which the ancestral population splits to form two daughter populations of the size specified in the panel. Each column is a separate set of simulations, with the top row plotting Z against tMRCA (measured in N generations, where N is the population size) for the larger daughter population, and the bottom row the smaller. Deleterious mutations are drawn from a gamma DFE with parameters inferred from *Drosophila melanogaster* population data.

**Supplementary Figure S7:**
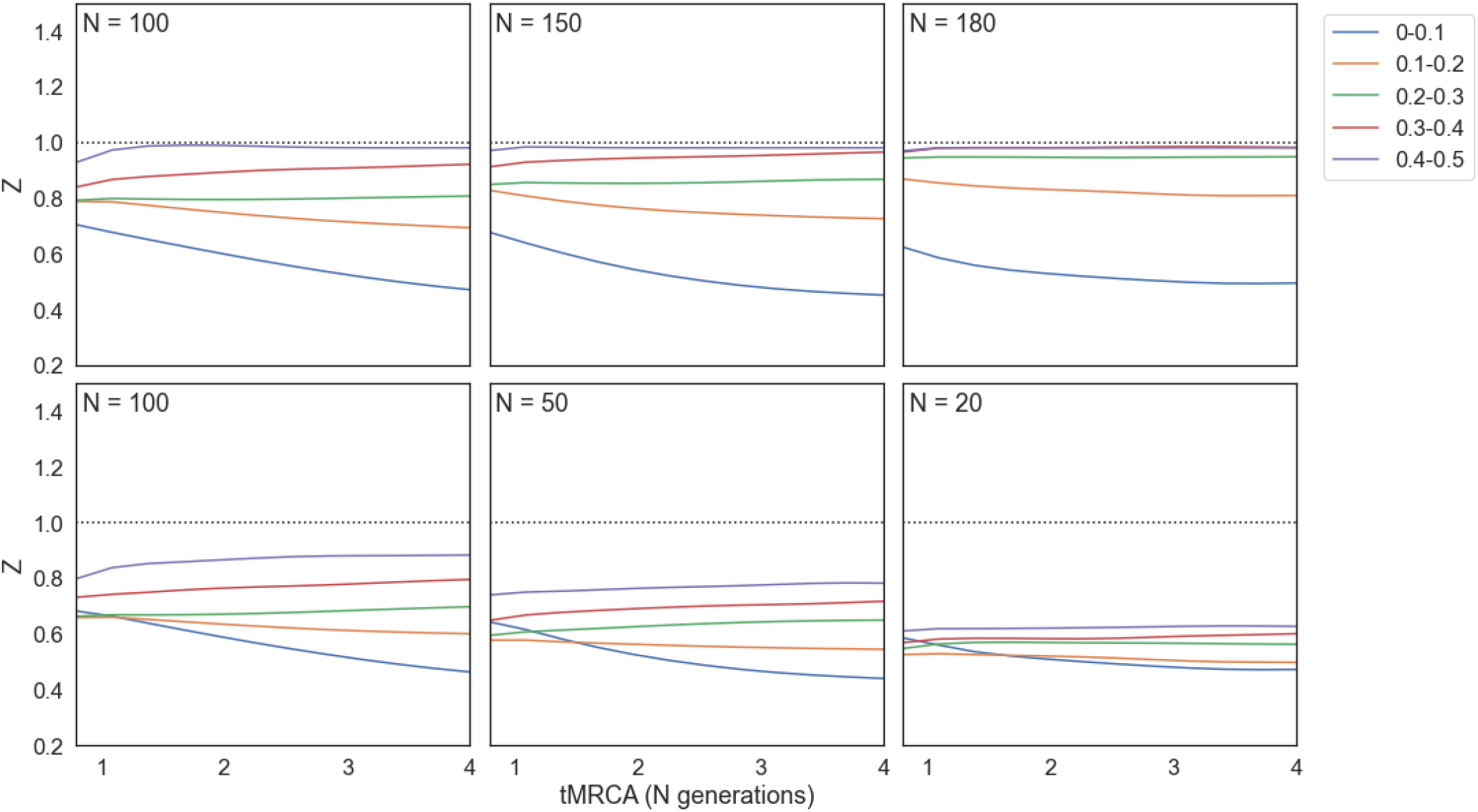
Dispersal simulations in which a single daughter population disperses from the ancestral population. Each column is a separate set of simulations, with the top row plotting Z against tMRCA (measured in N generations, where N is the population size) for the ancestral population, and the bottom row the daughter population. Deleterious mutations are drawn from a gamma DFE with parameters inferred from *Drosophila melanogaster* population data.

**Supplementary Figure S8:**
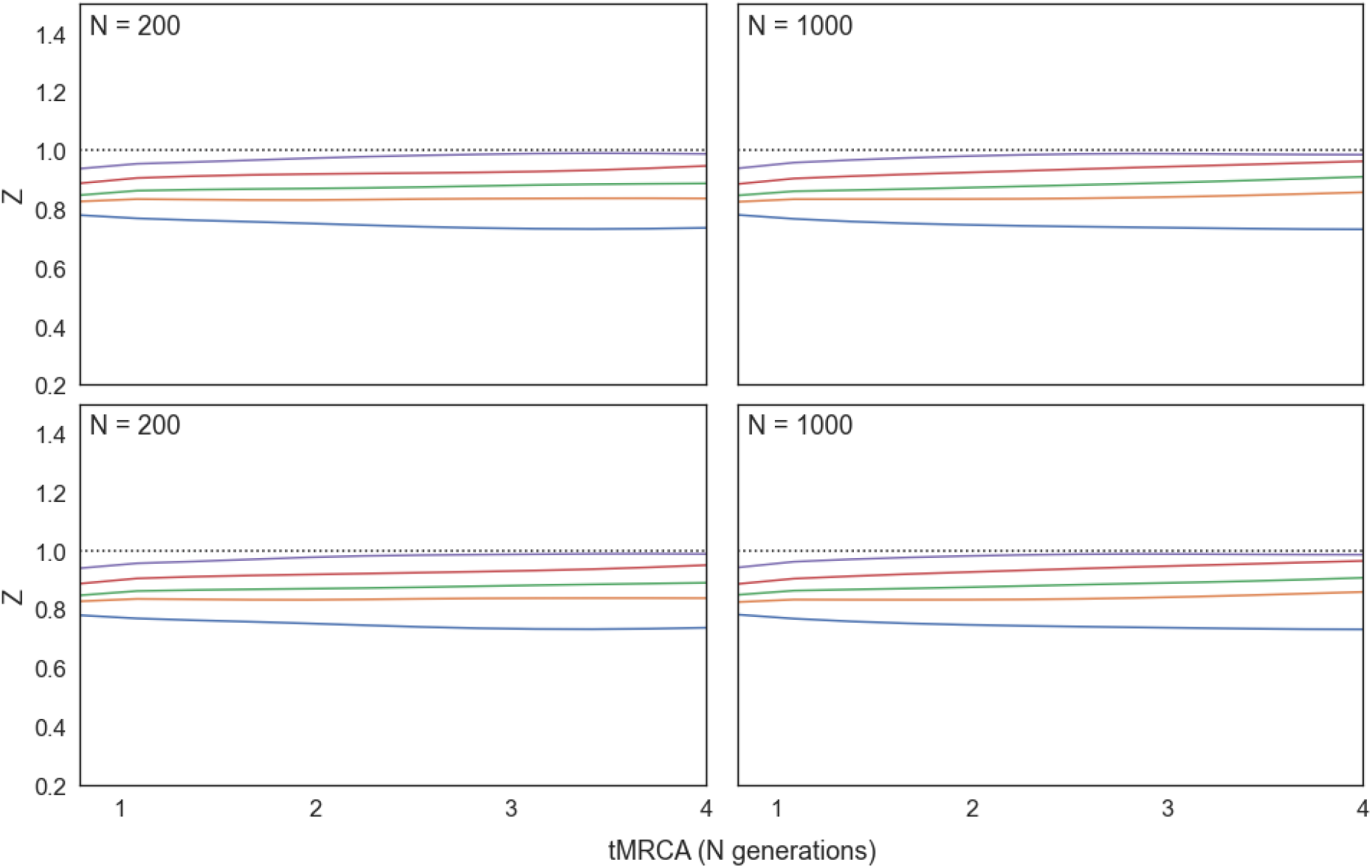
Vicariance expansion simulations in which both daughter populations expand. The ancestral population (of size N=200) splits to form two daughter populations of size N=100. Both daughter populations go on to expand in size. In the left column the daughter populations double in size. In the right panel they reach 10x their initial size. Deleterious mutations are drawn from a gamma DFE with parameters inferred from *Drosophila melanogaster* population data.

**Supplementary Figure S9:**
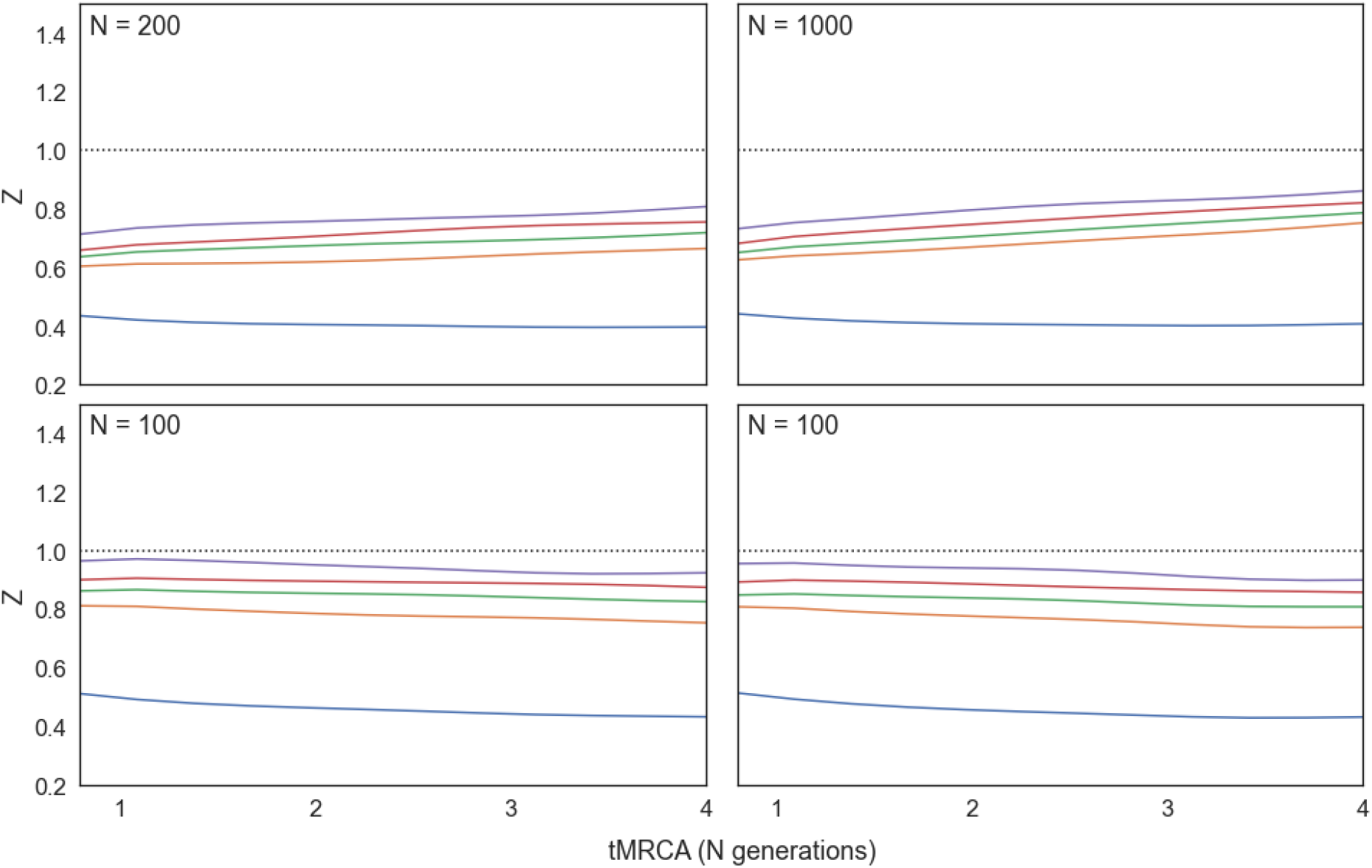
Vicariance expansion simulations in which only one daughter population expands. The ancestral population (of size N=200) splits to form two daughter populations of size N=100. One daughter population (upper panels) goes on to expand in size. In the left column the daughter populations double in size. In the right panel they reach 10x their initial size. Deleterious mutations are drawn from a gamma DFE with parameters inferred from *Drosophila melanogaster* population data.

**Supplementary Figure S10:**
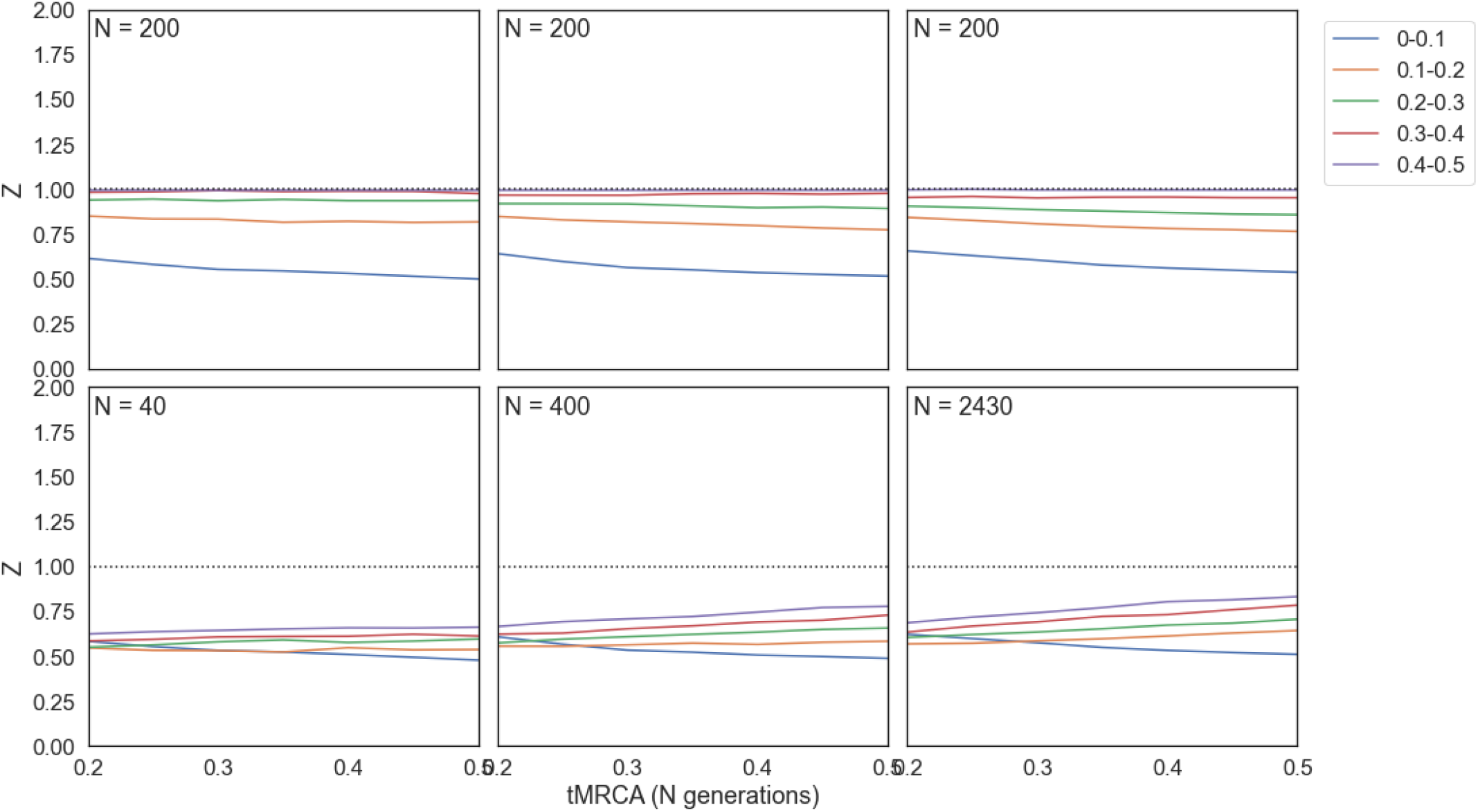
Dispersal expansion simulations in which a single daughter population disperses from the ancestral population and then expands. The ancestral population (of size N=200) splits to form a daughter population of size N=100, which expands to the final population size shown in the panel. Each column is a separate set of simulations, with the top row plotting Z against tMRCA (measured in N generations, where N is the population size) for the ancestral population, and the bottom row the daughter population. Deleterious mutations are drawn from a gamma DFE with parameters inferred from *Drosophila melanogaster* population data.

**Supplementary Figure S11:**
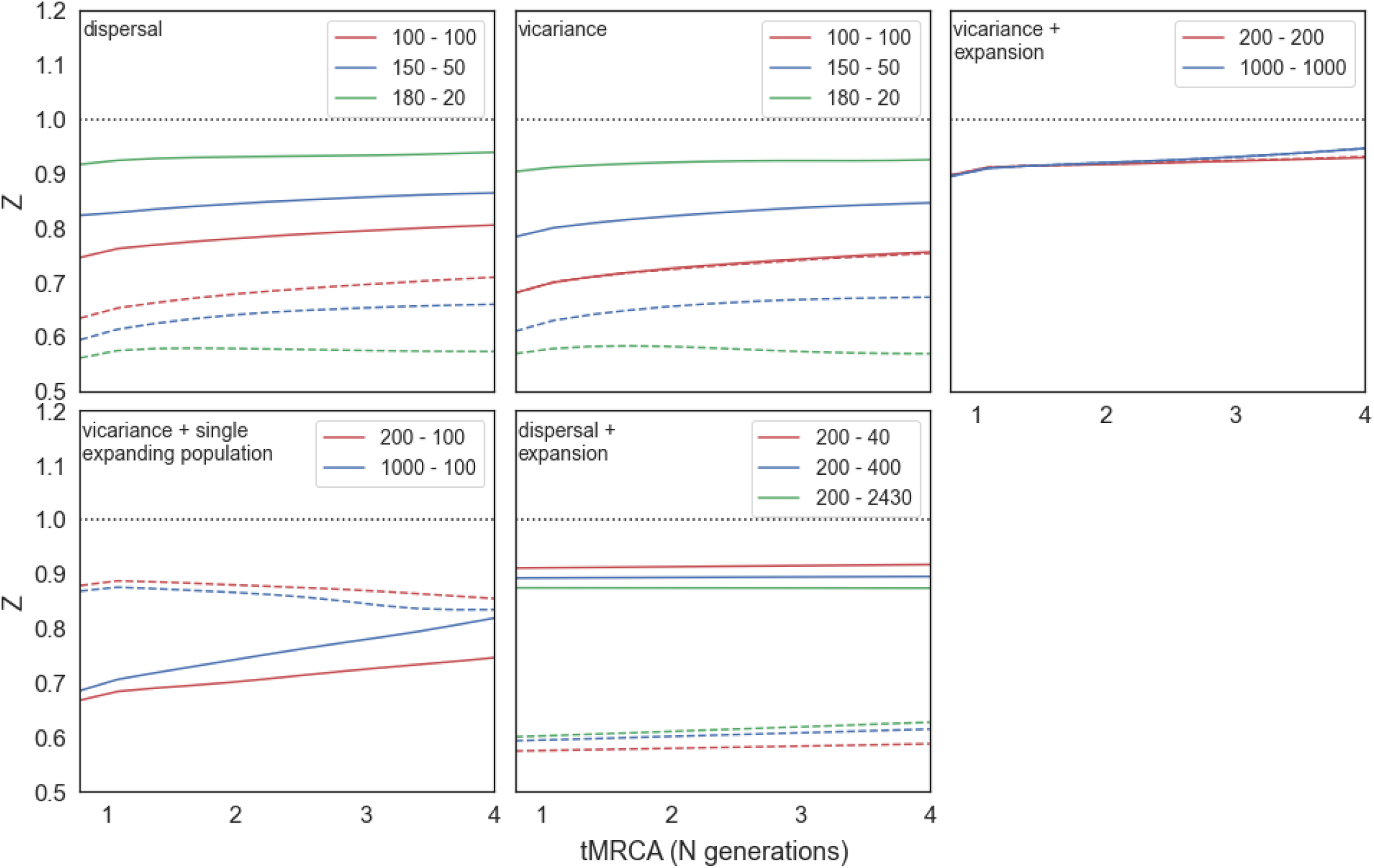
Simulations with for combined 0.1-0.5 minor allele frequencies. Each panel is a separate simulated scenario, with population sizes listed in the panel legend. The first number is for the filled in data lines, denoting the ancestral population in dispersal scenarios, and for the larger population in the vicariance scenarios. The second number is for the dotted data lines, denoting the daughter population in dispersal scenarios, and the smaller population in the vicariance scenarios. For more details on each scenario please see supplementary figures S1-10. Deleterious mutations are drawn from a gamma DFE with parameters inferred from human population data.

**Supplementary Figure S12:**
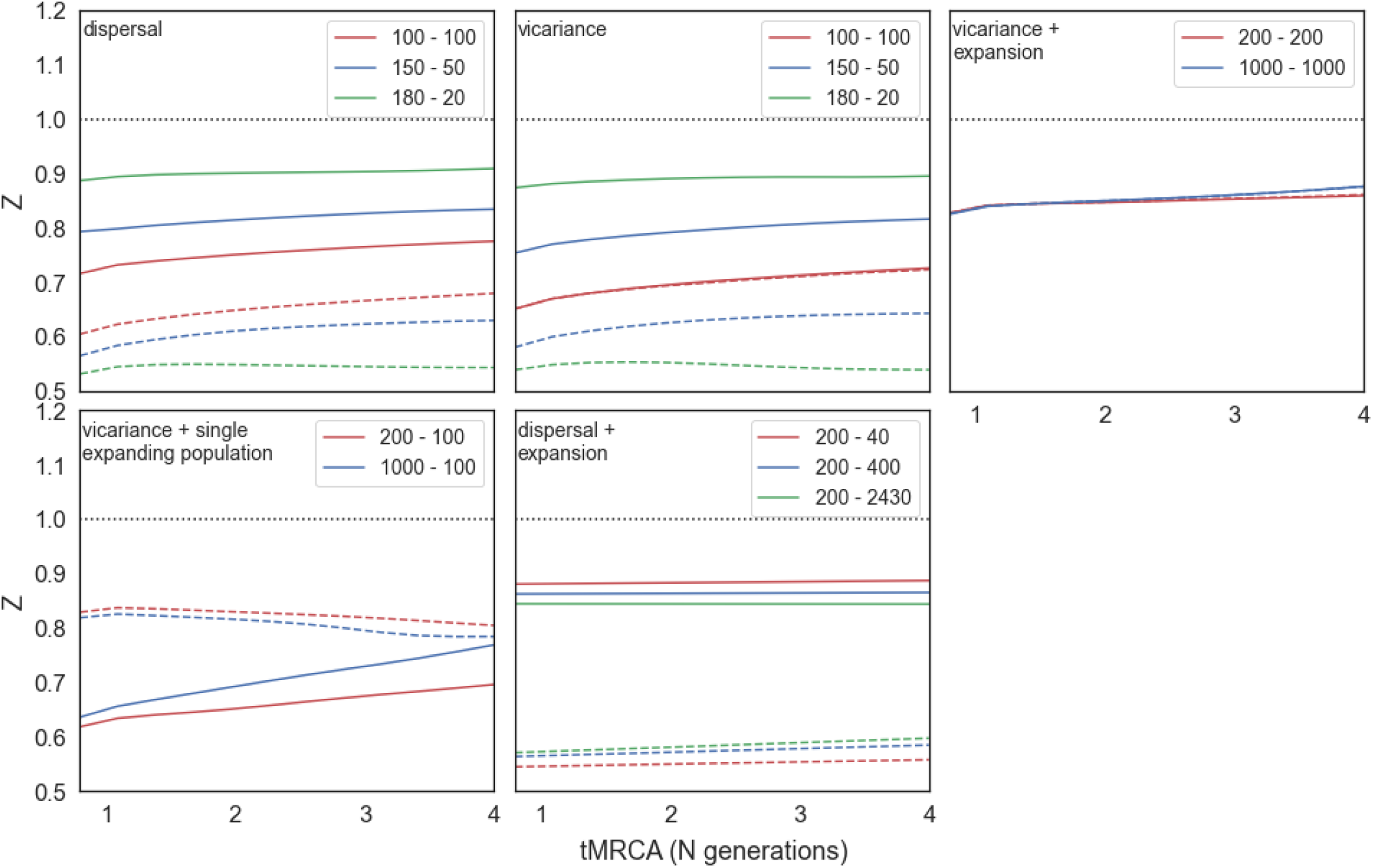
Simulations with for combined 0.1-0.5 minor allele frequencies. Each panel is a separate simulated scenario, with population sizes listed in the panel legend. The first number is for the filled in data lines, denoting the ancestral population in dispersal scenarios, and for the larger population in the vicariance scenarios. The second number is for the dotted data lines, denoting the daughter population in dispersal scenarios, and the smaller population in the vicariance scenarios. For more details on each scenario please see supplementary figures S1-10. Deleterious mutations are drawn from a gamma DFE with parameters inferred from *Drosophila melanogaster* population data.

**Supplementary Figure S13:**
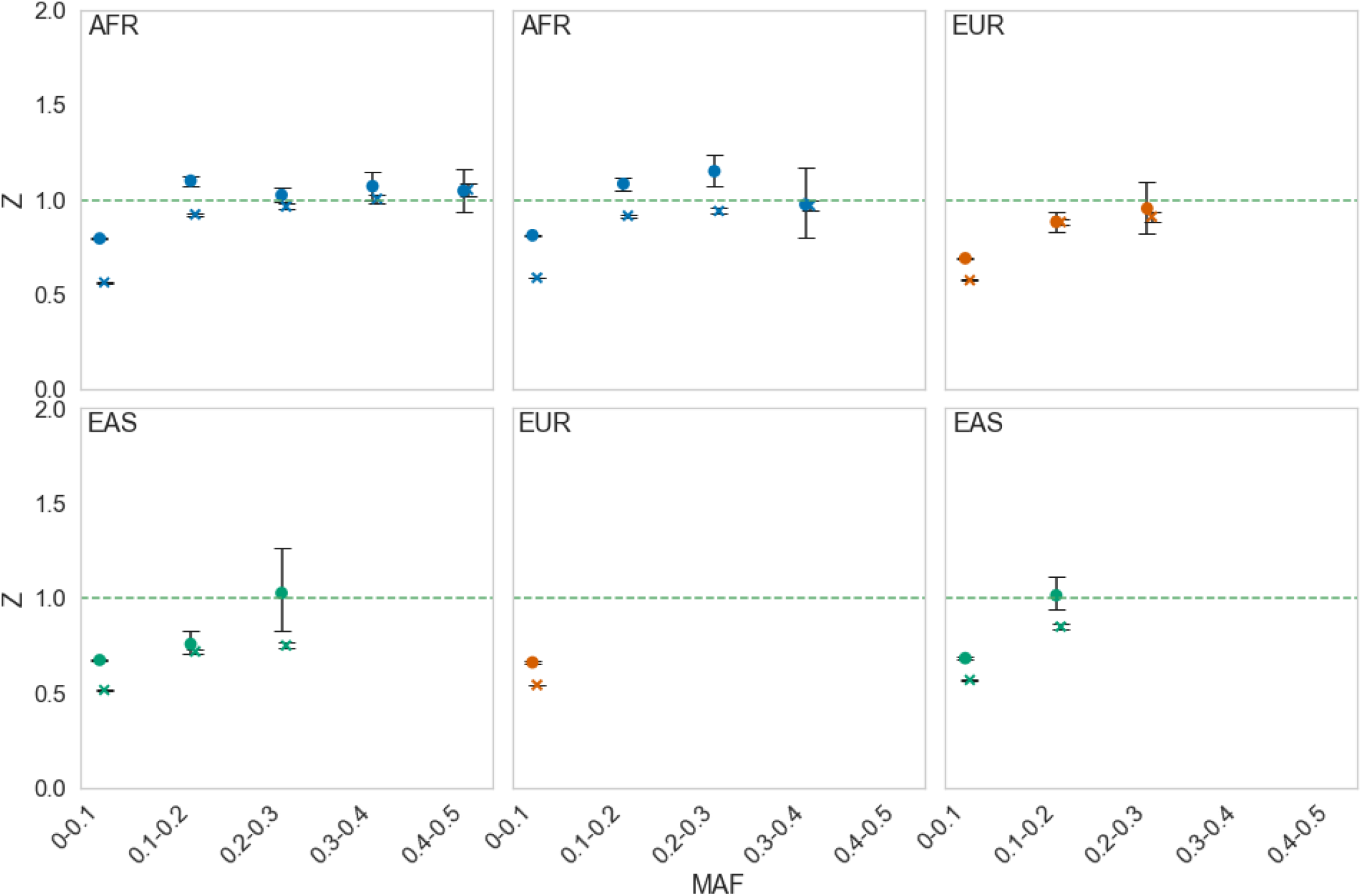
Simulations using the Gravel model of human demography (Gravel et al, 2011). Shown are the observed (filled circles) and simulated (crosses) values of Z. Each column represents a different population comparison. From left to right: Africans (AFR) and East Asians (EAS), Africans and Europeans (EUR), Europeans and East Asians. The population name in the upper left indicates which set of private polymorphisms are used to calculate Z in each population comparison. The x-axis represents private polymorphism minor allele frequency bins. Confidence intervals generated by bootstrapping.

**Supplementary Figure S14:**
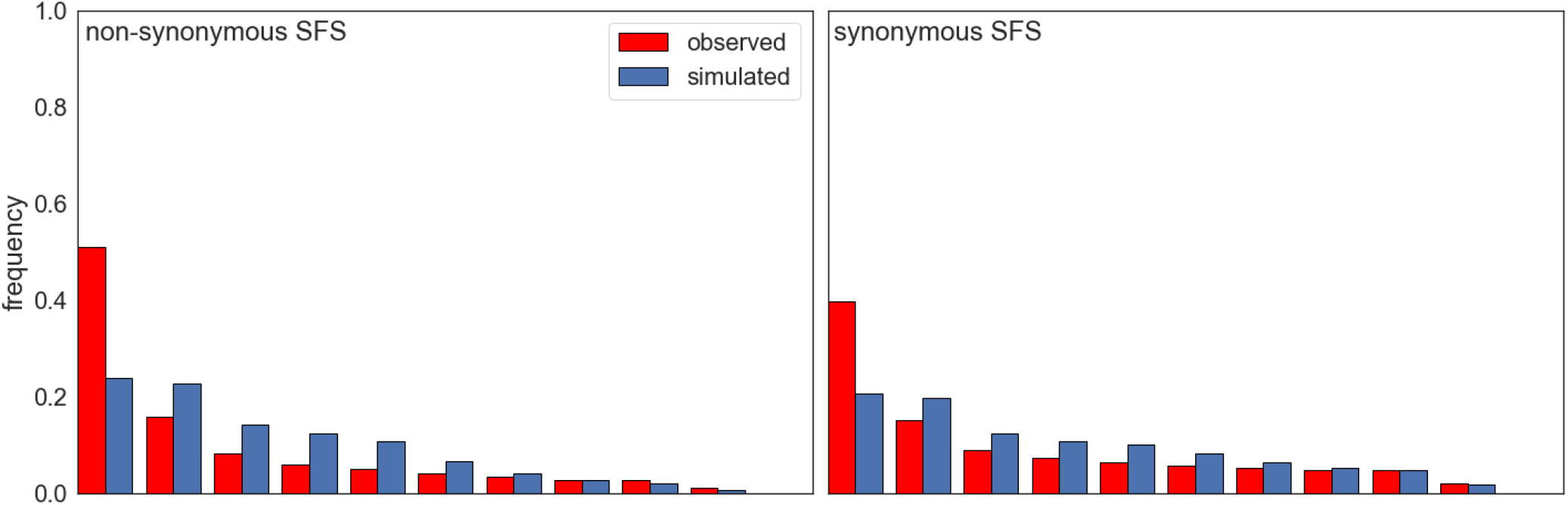
Comparison of simulated (under the Gravel et al. (2011) model of human demography) and observed SFS from the African population.

